# Cargo-mediated mechanisms reduce inter-motor mechanical interference, promote load-sharing and enhance processivity in teams of molecular motors

**DOI:** 10.1101/2021.06.10.447989

**Authors:** Niranjan Sarpangala, Ajay Gopinathan

## Abstract

In cells, multiple molecular motors work together as teams to carry cargoes such as vesicles and organelles over long distances to their destinations by stepping along a network of cytoskeletal filaments. How motors that typically mechanically interfere with each other, work as teams is unclear. Here we explored the possibility that purely physical mechanisms may potentially enhance teamwork, both at the single motor and cargo level. To explore these mechanisms, we developed a 3D dynamical simulation of cargo transport along microtubules by teams of canonically non-cooperative kinesin-1 motors. We accounted for cargo membrane fluidity by explicitly simulating the Brownian dynamics of motors on the cargo surface and considered both the load and ATP dependence of single motor functioning. We showed explicitly, for the first time, that surface fluidity leads to the reduction of negative mechanical interference between kinesins, enhancing load sharing thereby decreasing single motor off-rates and increasing processivity. Remarkably, we also showed that, independent of fluidity, increasing numbers of bound motors pull the cargo closer to the microtubule, increasing the on-rates of individual unbound motors, resulting in a cooperative increase in bound motor numbers that depends on 3D cargo geometry. At the cargo level, surface fluidity makes more motors available for binding, though this effect is significant only at low ATP or high motor density. Interestingly, we find that the fluidity induced reduction in mechanical interference dominates over the increased availability of motors for typical physiological and in vitro conditions, this allowing us to reconcile different experimental results in different regimes. Finally, we show that these effects can altogether result in enhanced mechanical efficiency and tunable cargo run-lengths for teams of molecular motors over physiological ranges of fluidity with implications for new experimental validation efforts, transport *in vivo*, artificial cargo design and multi-motor-driven mechanical processes in general.

**Author summary:** In cells, multiple molecular motors work together as teams to carry cargoes such as vesicles and organelles over long distances to their destinations by stepping along a network of protein filaments. How do intrinsically non-cooperative motors function as teams within cells? In this paper, we show, using computer simulations, that the fluid surfaces of cellular cargo reduce the mechanical interference between motors allowing better load sharing and decreasing their detachment rates, thereby increasing the distance over which they can carry cargo. We additionally found that motors pull cargo closer to the filament increasing the attachment rates of other unbound motors. These effects act synergistically with an increased availability of motors due to fluidity to further increase travel distances. Our simulation model accounted for the dynamics of the motors on the cargo surface and their load and ATP dependent kinetics, allowing us to connect single motor properties to overall transport. Our work on understanding how teamwork arises in mechanically coupled motors sheds new light on cellular processes, reconciles existing observations, encourages new experimental validation efforts and can also suggest new ways of improving mechanical efficiency and transport of artificial cargo powered by motor teams.

## Introduction

In eukaryotic cells, the intracellular transport of material between various organelles and the cellular and nuclear membranes is critical for cellular function and is an active process facilitated by the consumption of energy [1, 2]. A variety of motor proteins including myosins, kinesins and dyneins convert chemical energy from ATP hydrolysis into directed stepping along cytoskeletal protein filaments such as actin filaments and microtubules [1, 3]. These motors move in different directions along the filaments typically carrying lipid bilayer vesicles packed with proteins and signaling molecules or even membrane bound organelles such as Golgi and mitochondria [1, 2]. Defects in motor function can impair normal cell functioning and lead to a variety of pathologies including neurodegenerative diseases like Alzheimer’s disease and ALS [4–9]. Given its importance *in vivo*, there has been a lot of work done over the years in understanding motor function in the transport context ranging from single molecule studies that elucidate the detailed mechanistic workings of motors [10–17] to the functioning of multiple motors and their co-ordination [18–27]. Several theoretical and *in vitro* experimental studies [28–32] indicate that the collective behavior of motors is critically influenced by their coupling to each other resulting in observable effects in transport speeds and run lengths. For example, it is known that non-cooperative kinesin motors coupled together by a rigid cargo interfere with each other’s functioning leading to enhanced detachments and lowered run lengths [28–30]. How then are *in vivo* cargoes such as membrane-bound vesicles and organelles typically carried over long distances by teams of kinesin motors [33, 34]? It is possible that a fluid membrane simply makes more motors available for binding at the filament, thereby enhancing processivity [35, 36]. Alternatively or in addition, there could be other coupling-dependent physical mechanisms that directly affect single motor functioning and thereby promote teamwork. Studies on transport *in vivo* could address these questions directly, but the environmental complexity makes it difficult to disentangle fundamental phenomena of physical origin from the effects of other regulatory mechanisms [37]. *In vitro* systems, which offer cleaner insights, include microtubule gliding assays on flat bilayers with embedded motors [35, 38, 39], motor driven nanotube extraction from vesicles [40] and more recently even membrane covered beads carried by kinesin motors [41]. Studies of these systems and related models have generated a variety of interesting and sometimes conflicting results on cargo run lengths and velocities in the presence of membrane fluidity [36, 38, 41, 42]. These results are also obtained under different conditions of cargo geometry, motor density, cargo surface fluidity and environmental factors like ATP concentration. In order to reconcile them, we need an understanding of the relative importance and interplay of these factors and their effects on the number of engaged motors and the loads they experience, which in turn influences their collective speed and processivity. More importantly, to uncover any physical mechanisms present in teams of motors that directly affect single motor functioning and thereby enhance teamwork, we need to be able to monitor both single motor dynamics and collective cargo transport simultaneously.

Here, we used Brownian dynamics simulations to accomplish this goal. Briefly, we model the transport of a spherical cargo with a fluid surface carried by multiple kinesin-1 motors along a microtubule (MT) (see Fig. 1 and details in Materials and Methods). We explicitly accounted for the movement of the attachment points of the motors on the cargo surface by allowing free diffusion for unbound motors and diffusion biased by exerted forces for bound motors, with cargo surface fluidity determining the diffusion constant. We also accounted for the force and ATP dependence of the bound motor’s off-rate and stepping rate. The spherical cargo’s position was then updated by Brownian dynamics depending on the net force exerted by all bound motors. Using this model enabled us to focus on the dynamics of the motors as they diffused on the surface, their binding and unbinding from the filament and the forces experienced by individual motors, as a function of motor diffusion constant, motor density and ATP concentration. We connected these characteristics at the individual motor level to the effect of surface fluidity (inverse viscosity proportional to the diffusion constant) on the collective behavior of motors at the cargo level by analyzing transport properties such as the average number of bound motors, speed and distance traveled (run length). We uncovered two salient features, the reduction of interference and the cooperative increase of on-rates. We showed explicitly, for the first time, that surface fluidity leads to the reduction of negative mechanical interference between kinesins, characterized by lower forces on individual motors and a significant drop in antagonistic forces between motors. This allows teams of fluid-coupled motors to more fully exploit load sharing without inter-motor interference. This decreases single motor off-rates and increases processivity. Interestingly, increasing numbers of bound motors pull the cargo closer to the microtubule, increasing the on-rates of unbound motors, resulting in a cooperative increase in bound motor numbers that depends on 3D cargo geometry. Surface fluidity also makes more motors available for binding, as expected, and as indicated by previous studies [35, 36, 42]. However, this effect, by itself, is significant only at lower ATP concentrations (and/or very high motor numbers) when the effective timescale for diffusion and binding is less than the unbinding time. In fact, we estimate that the reduction in interference is the most significant effect under physiological conditions of high ATP and low motor numbers. Taken altogether these effects cooperatively result in increased mechanical efficiency and processivity with an increase in fluidity for teams of molecular motors, with both technological and *in vivo* implications.

**Fig 1.**
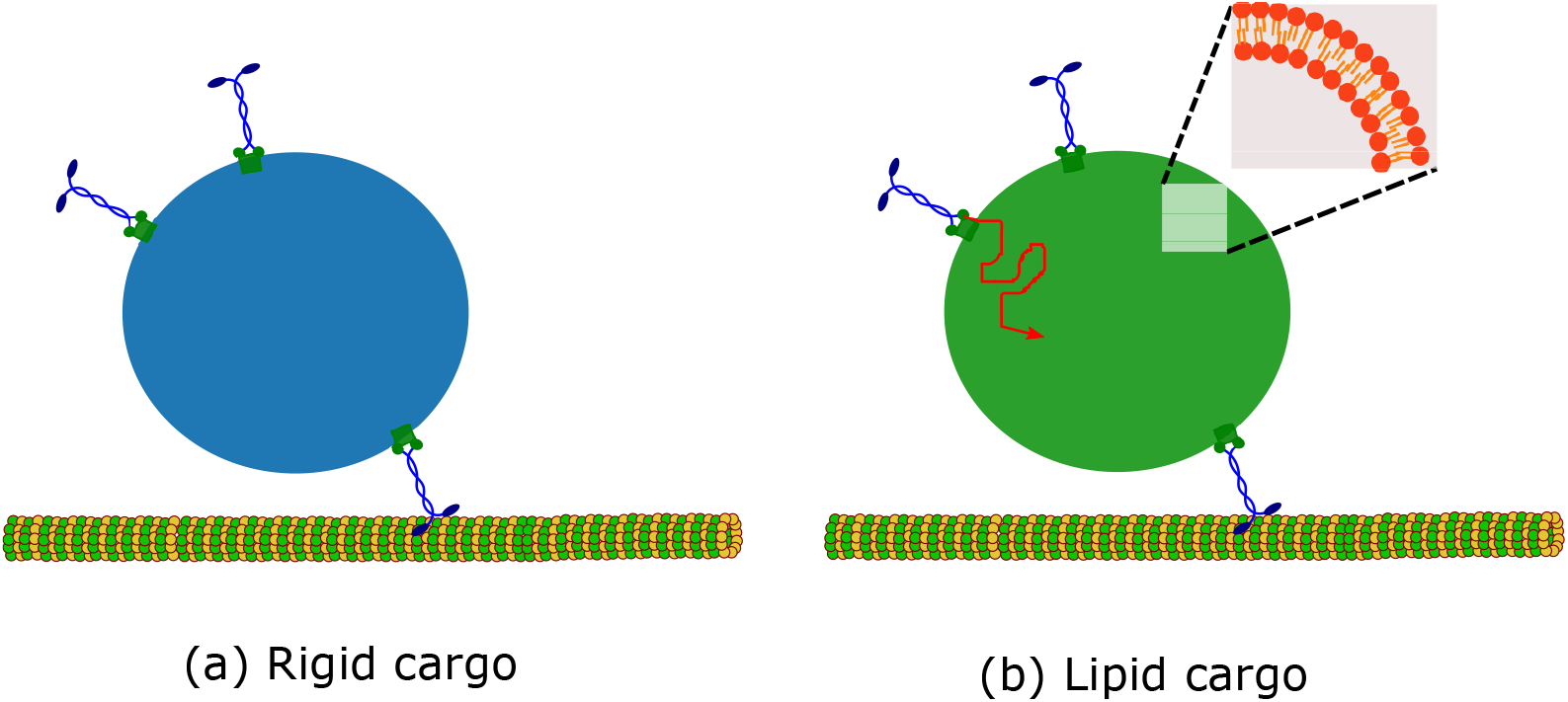
Two kinds of cargoes considered in the study. (a) Rigid cargo (membrane-free cargo). Molecular motors are permanently attached to random locations on the cargo surface. (b) Fluid/Lipid cargo (membrane-enclosed cargo). Molecular motors can diffuse on the cargo surface. Inset of Fig. 1(b) The lipid bilayer that forms the fluid cargo surface.

## Results

### Surface fluidity reduces negative interference between bound kinesin motors

We first addressed the question of whether the fluidity of the cargo surface has any influence on the functioning of a bound kinesin motor. We expected that the motor attachment point’s freedom to move on the lipid cargo surface in response to forces could relieve strain and reduce the forces experienced by the motor. Fig. 2(a) shows a visual comparison of the time traces of the individual motor forces experienced by a team of three bound motors carrying a cargo in the rigid and lipid cases. The noticeably lower magnitudes of the peak forces experienced by the motors in the lipid cases suggest that our expectation is valid. To explore the effect more quantitatively, we computed the distribution of forces experienced by a single motor when it is part of a team of *n* motors that are simultaneously bound to the microtubule. This distribution, shown for *n* = 3 bound motors in Fig. 2(b), is significantly broader, reflecting larger forces, for a motor on a rigid cargo compared to any of the lipid cargoes. The lipid cargo motor force distributions for different diffusion constants are, in fact, comparable to that for a single bound motor carrying a rigid cargo. A rigid coupling of motors, therefore, results in individual bound motors experiencing higher forces than when they are singly bound to cargo, indicating negative interference. The introduction of surface fluidity reduces this interference by effectively decoupling the motors, consistent with our expectations. This reduction is apparent in the distribution of forces experienced both against (hindering) and in (assistive) the direction of processive motion (S1 Fig). This reduction is also evident in the fraction of “high” forces with a magnitude ≥ 0.01*F*_*s*_ (S2 Fig), where *F*_*s*_ is the stall force of kinesin motor (Figs. S3 and S4 show the fractions in hindering and assistive directions). For rigid cargo, this fraction increases as a function of the number of bound motors, *n*, indicating negative interference between motors. In the presence of surface fluidity, however, no such increase in the high force fraction with *n* is apparent, indicating a significant reduction of negative interference. In fact, the fraction decreases with *n* at high surface fluidity (*D* = 1 *μm*^2^*s*^−1^), which is likely due to the decrease in the average force experienced by the motors in the presence of multiple bound motors (S5 Fig). While the decrease in average force occurs for rigid cargo as well, the increase with *n* due to negative interference dominates.

**Fig 2.**
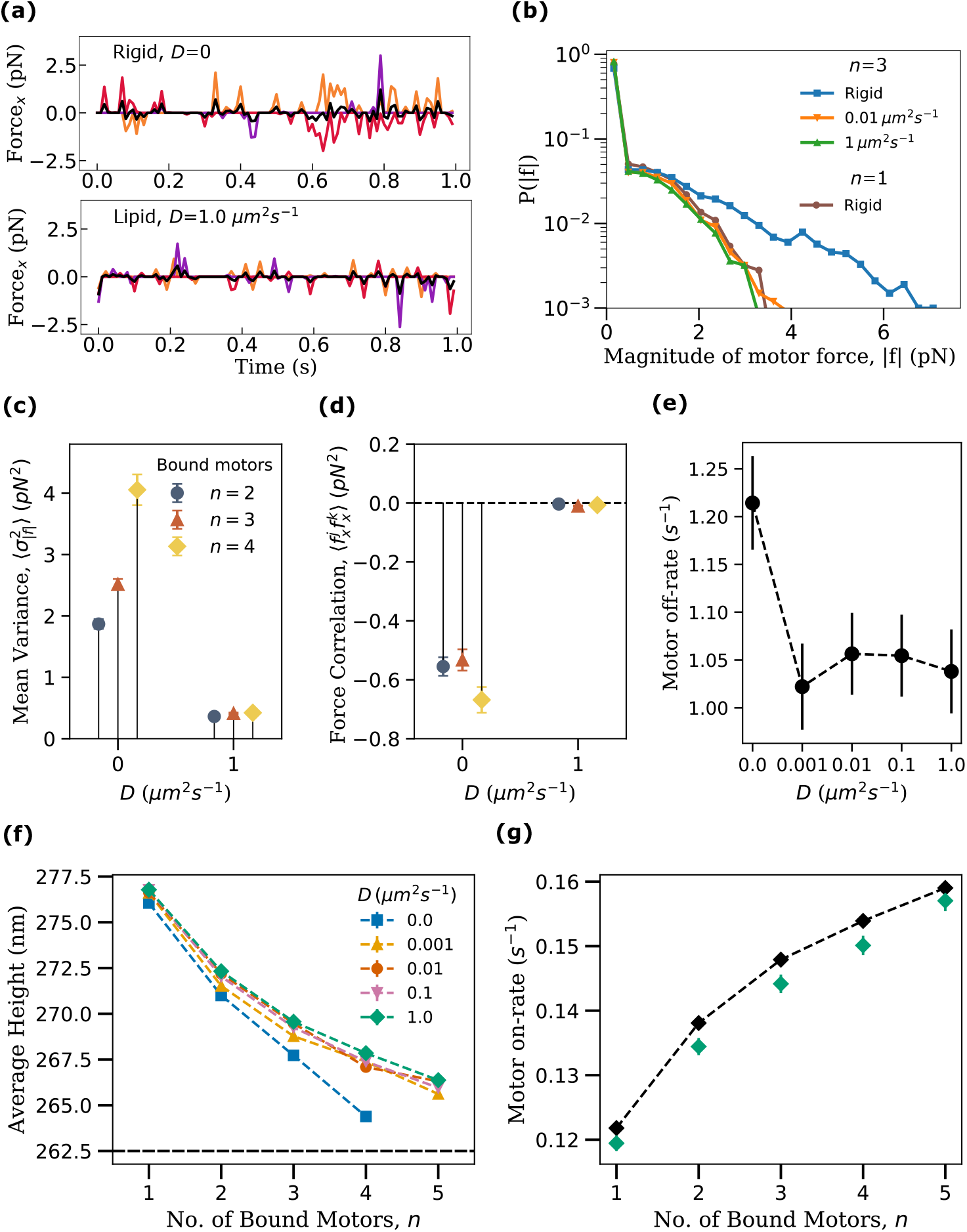
**(a)** The *x*-component of forces experienced by three simultaneously bound motors in rigid (top) and a lipid (bottom) cargoes. Colored curves indicate forces on individual motors, black curve represents the mean of all three. **(b)** Distributions of the magnitude of the force experienced by motors for different cargo surface fluidity at a fixed number of bound motors (*n* = 3). Distribution for singly bound motor carrying a rigid cargo is also shown for comparison. Sample size for each distribution curve was *S* = 10000 time points drawn from 117, 96 and 48 cargo runs for *D* = 0, 0.01, 1 *μm*^2^*s*^−1^ respectively. **(c)** The mean variance of forces among bound motors calculated as 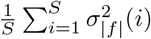 where the summation is over all the sample time points when the number of bound motors is *n*. 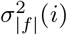 is the variance in the magnitude of force experienced by the *n* bound motors at *i*^*th*^ sample. Sample size *S* = 10000 drawn from 200 cargo runs. **(d)** Mean value of the correlation between the *x*-components of motor forces, 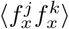, averaged over motor pairs and time points. Sample size *S* = 10000 drawn from 200 cargo runs. **(e)** Average value of motor off-rate as a function of fluidity of cargo surface (*S* = 580 from 200 cargo runs). **(f)** Average measured distance (h) of the center of mass of cargo from MT as a function of the number of bound motors and fluidity. Horizontal line indicates the height below which the cargo experiences steric interaction from the MT. Data for (a-f) were obtained from the simulation of transport of cargoes with *N* = 16 motors at high ATP concentration of 2 mM. Data sampling rate was 100 *s*^−1^. **(g)** Computational measurements of single motor on-rate (green diamonds) and analytical approximation (black dashed line with diamond). (*D* = 1 *μm*^2^*s*^−1^, *S* = 10000) using the cargo height for a given *n* from the data in (f). Error bars for all plots represent the standard error of the mean.

The variance between the individual forces experienced by simultaneously bound motors is also a measure of negative interference as it quantifies the deviation from perfect load-sharing where the variance is zero. The mean variance over all sampled instances with *n* bound motors shows a dramatic decrease at higher fluidity (*D* = 1 *μm*^2^*s*^−1^) compared to the rigid cargo case (Fig. 2(c)). In fact, even the presence of a small amount of fluidity accounts for most of the significant drop in variance (S6 Fig) indicating that fairly good load-sharing can be achieved with modest fluidity. We also quantified negative interference more directly by studying whether individual motors were acting antagonistically, i.e., exerting forces in opposite directions. To do this, we computed the correlation between the x-components of bound motor forces 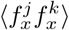 averaged over different motor pairs (*j, k*) and time (Fig. 2(d)). We see that this correlation is highly negative for rigid cargo indicating opposing motor forces while it is negligible in the presence of fluidity indicating motors on the same side of the cargo pulling in the same direction. Taken together, our results show that interference dominates over load sharing for rigid cargo while increasing surface fluidity reduces this interference and promotes load-sharing.

Next, we addressed the implications of the reduction in negative interference for the functioning of motors in the context of transport. The off-rate of a kinesin motor increases with the magnitude of the force that it experiences and also depends on its direction [1, 2, 15, 23, 43]. The observation that surface fluidity reduces negative interference leads us to expect that this will also lead to lower off-rates for individual motors. We measured the mean off-rate of motors as the inverse of the mean time spent by individual motors between a binding and subsequent unbinding event. We found that bound motors on fluid cargoes do indeed have lower off-rates than those on rigid cargoes (Fig 2(e)). Consistent with our findings for the forces and interference, the introduction of a small amount of fluidity accounts for most of the significant drop in off-rates with no significant variation with increasing diffusivity. Thus, increasing surface fluidity reduces inter-motor interference and promotes load-sharing leading to reduced off-rates and potentially longer processive runs for individual motors.

### On-rate of a motor increases with the number of bound motors for fluid cargo

The run length of a cargo carried by a team of motors depends on the interplay between the off-rates and on-rates of individual motors [23]. The on-rate of a motor, typically considered to be independent of the number of other bound motors, depends on its intrinsic binding rate when it is within reach of the MT. The region of the cargo surface (or access area) from where an anchored kinesin can reach the MT is defined, in our model, by the set of points on the surface at a distance less than or equal to the rest length of the motor (*L*_*mot*_ = 57 nm). For a cargo bound to the MT with one or more motors, this access area is a function of the cargo geometry and the proximity of the cargo to the MT [1, 35]. For rigid cargo, motors can only bind if they happen to be anchored in the access area while for fluid cargo, motors can diffuse in and out of the access area. For a given motor type with a fixed intrinsic binding rate and a given cargo geometry (a sphere here), the access area is then simply a function of the distance of the cargo from the MT. Interestingly, our measurements of the average distance of the center of mass of the cargo from the MT indicated that the cargo comes closer to the microtubule as the number of bound motors increases (Fig. 2(f)). This decrease in distance arises from the increase in the net vertical component of the average force due to an increased number of motors which serves to counteract cargo fluctuations away from the MT. We also note that cargoes with low surface fluidity are closer to the MT than the cargoes with high surface fluidity, especially for high numbers of bound motors. This is because motors on fluid cargoes can relax their tension by sliding on the cargo surface allowing for larger fluctuations (S7 Fig and S8 Fig) and therefore a higher average distance from the MT.

Thus, for both rigid and fluid cargo, as the number of bound motors increases, the cargo approaches the MT leading to an increase in the access area. We expect that this should lead to an increase in the on-rate of an individual unbound motor because its likelihood of being in the access area is increased. We verified this by measuring the on-rate of a motor diffusing on the surface of a cargo held at specific distances from the MT (Fig. 2(g)) that correspond to different numbers of bound motors (from data in Fig. 2(f) for *D* = 1 *μm*^2^*s*^−1^). Here we chose a high diffusion constant for simplicity and measured the mean time required for a randomly placed motor to diffuse and bind to the MT. Indeed, we see that the effective on-rate of the motor, calculated as the inverse of this mean time, increases with decreasing distance corresponding to increasing numbers of bound motors (Fig. 2(g)). To show that the quantitative increase in on-rate was explained by the decreasing distance, we also analytically estimated the change in on-rate. Since the average time spent by a motor in the access area before binding to the microtubule is approximately proportional to the ratio of access area to total area in the high diffusion constant limit, we estimated that the effective on-rate is *π*_*ad*_ = *π*_0_(*S*_*a*_*/S*_*T*_), where *π*_0_ = 5 *s*^−1^ is the intrinsic binding rate, *S*_*a*_ is the access area and *S*_*T*_ = 4*πR*^2^ is the total surface area of the cargo. By computing the access area numerically for different cargo distances, we were able to estimate *π*_*ad*_ corresponding to different numbers of bound motors. The rather good agreement with the measured values from simulations (Fig. 2(g)) indicates that the increase in on-rate is captured by the effects of increased access area. It is to be noted that the rate at which the cargo transitions from being bound by *n* to *n* + 1 motors can be calculated from this on-rate, the off-rate and the number of unbound motors available for binding (details in S2 Appendix and data in S9 Fig.)

Our results reveal a positive feedback effect at play. As the number of bound motors increases, the cargo gets closer to the MT and the on-rates of each of the remaining motors increase which further increases the number of bound motors. But a cargo cannot benefit from this positive feedback unless it makes a sufficient number of motors available for binding.

### Surface fluidity increases the availability of motors for binding

We expect that the diffusion of motors on the cargo surface should lead to a greater availability of motors in the access region and hence a higher number of bound motors compared to rigid cargo with the same overall number of motors. We also expect that this will be true only if *τ*_*bind*_ - the typical time for any one of the unbound motors to bind to the microtubule is less than *τ*_*off*_ = 1*/ϵ*_0_ - the typical time for a given bound motor to detach. The relative values of these two timescales are set by parameters such as the number of motors (*N*), the radius of cargo (*R*), and the ATP concentration. For instance, raising the motor density (say by raising *N*) will result in a higher effective on-rate and a decreased *τ*_*bind*_, while decreasing the ATP concentration, lowers the off-rate and raises *τ*_*off*_.

We first looked at a parameter set (*N*=16 [ATP]= 2 mM) satisfying the condition *τ*_*bind*_ < *τ*_*off*_ (see S2 Appendix for details of estimates). Fig. 3(a) shows that, while the average number of bound motors, 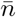, increases with time for both rigid and fluid cargoes, the fluid cargo indeed accumulates more motors. The spatial distribution of motors in Fig. 3(b) further highlights the difference between rigid and fluid cargoes with the probability of finding motors near the microtubule on a fluid cargo being significantly larger at late times than for rigid cargo. It is to be noted that the slight increase at late times for rigid cargoes is simply due to the higher contribution to the average of cargoes, that happen to have motors clustered together, surviving for longer times.

**Fig 3.**
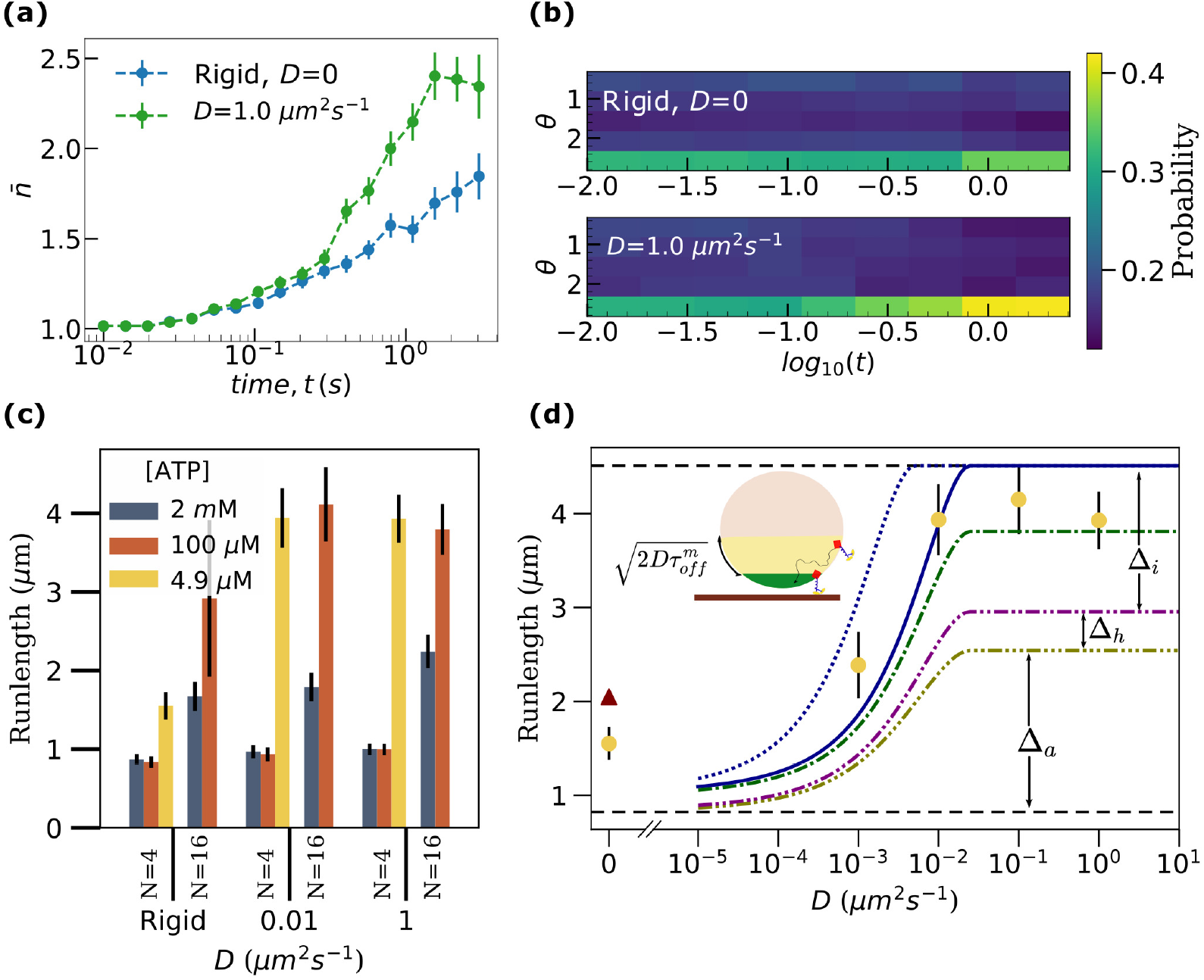
**(a)** Ensemble average of the number of bound motors, 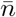 as a function of time (s). Error bars represent the standard error of the mean. **(b)** Spatial distribution of the kinesin motors on cargo, computed as the probability of locating motors as a function of the polar angle *θ* and time, *t. θ* is the polar angle of the motor, measured with respect to *z* axis perpendicular to the microtubule and passing through the center of the cargo. Data for Fig (a) and (b) was obtained from the transport of cargoes with a total of *N* = 16 motors at [ATP]=2 mM. **(c)** Runlength as a function of diffusivity (*D*) for two different numbers of motors on the cargo (*N*) and three different ATP concentrations. 200 cargo runs were considered for each parameter set. Error bars represent the standard error of the mean. **(d)** Comparison between run lengths from our simulations (*N* = 4, [ATP] = 4.9 *μM*, yellow circles) and analytical estimates (maroon triangle, solid, dashed and dash-dotted lines) as described in the text. *Inset cartoon*: A lipid cargo, with access area (*S*_*a*_, green) and influx area (*S*_*I*_, yellow) shown. Influx area is defined as the region within 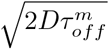 from the access area.

To further test the dependence of motor accumulation on the relative values of the two time scales, we considered four other sets of physiologically relevant parameters of *N* and [ATP] that had different relative magnitudes of *τ*_*off*_ and *τ*_*bind*_ (shown in Table 1) As predicted, there is no statistically significant difference in the average number of bound motors as a function of diffusivity (see S11 Fig(a)) for cases when *τ*_*off*_ ≈ *τ*_*bind*_ (*N* = 4, [ATP]= 2 *m*M and 100 *μ*M). The number of bound motors, however, increases as a function of diffusion constant (*D*) for cases where *τ*_*bind*_ < *τ*_*off*_ (*N* = 16 [ATP]= 2 *m*M and 100 *μ*M, *N* = 4 [ATP]= 4.9 *μ*M) as expected. This supports our expectation that fluid cargoes can indeed accumulate higher number of bound motors than rigid cargoes, but only under conditions that ensure that *τ*_*bind*_ < *τ*_*off*_ i.e. when the number of motors (*N*) is high or the ATP concentration is low or both.

**Table 1.**
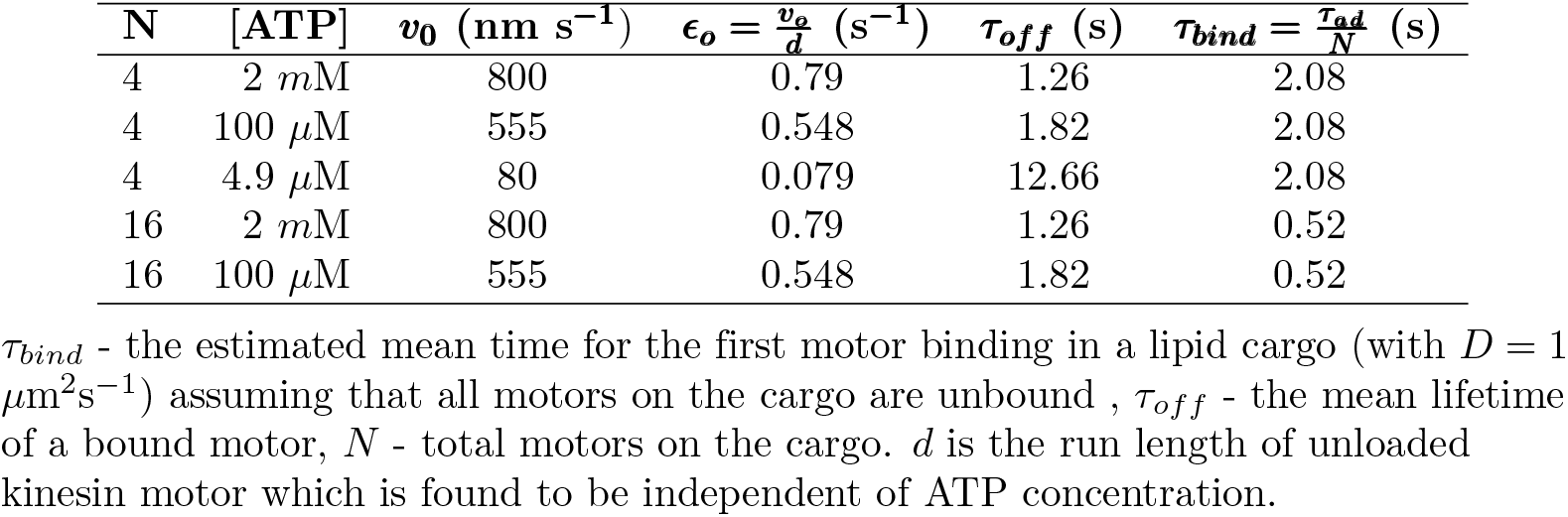
Comparison of the single motor binding and unbinding times for different N and [ATP].

### Fluid cargo have longer run lengths than rigid cargo

So far we have shown that increasing surface fluidity reduces inter-motor interference and promotes load-sharing leading to reduced off-rates. Fluidity also makes more motors available for binding in a cooperative fashion with increased binding leading to higher on-rates. We would expect that having more motors bind that survive longer should lead to increased run lengths for fluid cargo. The run lengths of cargoes measured as a function of diffusion constants (Fig. 3(c)) suggest that this is indeed possible given the right ATP concentration and the number of motors on the cargo (see S12 Fig for additional diffusion constants). Although the run length is unaffected by fluidity when *τ*_*bind*_ is of the order of *τ*_*off*_ (low *N* and medium/high [ATP]), the run length increases with the fluidity of the cargo surface when *τ*_*off*_ > *τ*_*bind*_ (for high *N* or low [ATP] or both). The effect is much more pronounced when the disparity between the timescales is higher.

We saw earlier that the average number of bound motors shares similar trends (S11 Fig) as a function of cargo surface fluidity. It is to be noted, however, that cargoes can have the same average number of bound motors but different run lengths. For example, although high fluidity (D= 1 *μm*^2^*s*^−1^) cargoes with *N* = 4 at [ATP]= 4.9 *μ*M and *N* = 16 at [ATP]= 2 mM have the same average number of bound motors, the lower ATP case with fewer total motors shows a substantially larger run length (S11 Fig(a) and S11 Fig(b)). This is a consequence of the higher tendency of a cargo to survive a 1 motor state at low ATP due to a lower motor detachment rate (more details in S4 Appendix). Overall, our results indicate that surface fluidity increases run lengths when the number of motors (*N*) is high or the ATP concentration is low or both.

We now consider the relative contribution to the overall run length of the different effects we showed are at play; the reduction in negative interference, the cooperative increase in on-rates and the increased availability of motors.

To estimate the magnitude of these contributions, we first consider an analytical expression for the run length [23]

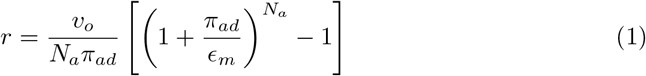

where *ν*_*o*_, *N*_*a*_, *π*_*ad*_, *ϵ*_*m*_ are the motor speed, the available number of motors for transport, the single motor binding rate and the mean motor off-rate respectively. Based on our results, we now generalize the expression by incorporating the dependence of these parameters on the cargo surface diffusivity *D* and the number of bound motors. The mean off-rate is sensitive to the presence of inter-motor interference and the introduction of fluidity almost completely eliminates interference (Fig. 2(b-e)). This means we can treat the mean off-rate, *ϵ*_*m*_, as having two values; a higher one for rigid cargo due to interference and a lower one for fluid cargo (Fig. 2(e)). The number of available motors, *N*_*a*_, is usually taken to be the number of motors in the access area, *S*_*a*_, for rigid cargo. For *D* > 0, however, additional motors from an influx area *S*_*I*_ (see Fig. 3(d)) can reach the access area before the cargo unbinds from the microtubule and contribute to transport. Here we estimate *S*_*I*_ as defined by a region within a distance 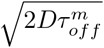 of the access area, representing the distance diffused by a motor over the mean motor lifetime 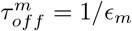. We, therefore, take *N*_*a*_ to be the average number of motors in the access and influx areas combined, *N*_*a*_ = 1 + (*N* − 1)(*S*_*I*_ + *S*_*a*_)*/S*_*T*_. Thus the number of available motors starts from a minimum value *N*_*a*_ = 1 + (*N* − 1)(*S*_*a*_*/S*_*T*_) at *D* = 0 and increases monotonically as function of *D* before saturating at *N*_*a*_ = *N*. Finally, motors bind with a rate *π*_*ad*_ = *π*_0_ = 5 *s*^−1^ when they are within the access area and do not bind otherwise. This means we can take the mean on-rate of the available motors to be *π*_*ad*_ = *π*_0_*S*_*a*_*/*(*S*_*I*_ + *S*_*a*_). We note that the access area, *S*_*a*_, depends on cargo geometry and in particular on the height of the cargo above the microtubule. As more motors bind, the height decreases, *S*_*a*_ increases and the effective on-rate increases (Fig. 2(f-g)). Finally, we note that the cargo velocity did not change by more than 5% over the range of parameters we tested (S14 Fig) and so we took *v*_*o*_ to be constant.

Using these analytic estimates, we can compute approximate bounds for the relative contributions of the different effects to overall run length. As an example, we do this for the case of low motor density and low ATP (*N* = 4 [ATP]= 4.9 *μ*M, Fig. 3(d), other cases in S16 Fig). We start with the limiting lower bound case where there is no additional availability of motors due to diffusion (*N*_*a*_ = 1 + (*N* − 1)(*S*_*a*_*/S*_*T*_)), no reduction in interference (*ϵ*_*m*_ = 0.12 *s*^−1^, S15 Fig) and the cargo height is at its maximum (corresponding to having 1 motor bound (Fig. 2(f))). This yields the horizontal dashed line at the bottom in Fig. 3(d). A similar upper bound that assumes all motors are available (*N*_*a*_ = *N*), interference is absent (*ϵ*_*m*_ = 0.1 *s*^−1^, see S15 Fig) and the cargo height corresponds to that at *n* = 2 (which is close to the average number of bound motors for this configuration), yields the horizontal dashed line at the top. While the bounds bracket the run lengths measured from the simulations, we notice that the lower limit is well below the actual values, especially at higher diffusion constants. It is to be noted that the bounds do not really extrapolate well to *D* = 0 because, for rigid cargo, the average run length is not set by the average number of motors in access area. A better approximation is the weighted average of run lengths for different numbers of motors in the access area weighted by the probability of having that many motors in the access area (maroon triangle in Fig. 3(d); details in S3 Appendix).

We first consider the contribution due to diffusion increasing the availability of motors alone. Allowing *N*_*a*_ to increase with *D* but with no reduction in interference (*ϵ*_*m*_ = 0.12 *s*^−1^) and the maximum cargo height, corresponding to *n* = 1, yields the lowest dash-dotted curve in Fig. 3(d). While there is a substantial increase in run length with increasing *D*, this increase (of order Δ_*a*_ at high *D*) yields an estimate that is still significantly below the measured values at high *D*. This indicates that the increased availability of motors alone cannot account for the entire increase in run length. We next incorporate the cooperative increase in the accessible area due to more motors binding and the cargo moving closer to the microtubule by setting the cargo height to that corresponding to *n* = 2 bound motors. This yields an increase in the run length (middle dash-dotted curve in Fig. 3(d)) but the gain, of order Δ_*h*_ at high *D*, is also too small to account for the observed increase. We finally incorporate the effect of reduced interference between the motors due to surface fluidity by taking *ϵ*_*m*_ = 0.1 *s*^−1^. This shifts the run lengths upwards considerably, by Δ_*i*_ at high *D*, and brings the estimates to within the range of the observed values (solid curve in Fig. 3(d); the upper dash-dotted curve is for a cargo height corresponding to one bound motor). Intriguingly, the surface fluidity based reduction in negative interference has an effect that is comparable or even stronger (see S16 Fig also) than the effect due to the increase in the number of available motors. We note that a reduction in the interference and the resultant reduction in mean off-rate leads to an increase in the mean motor lifetime, *τ*_*m*_, and hence an increase in the influx area *S*_*I*_ which is set by 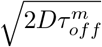. Thus, reduction in interference contributes to an increased run length not only due to an increased lifetime of each motor but also due to the resultant increase in the number of motors available for transport. While we used the single motor lifetime 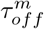 to estimate *S*_*I*_, this represents a lower bound since new motors may reach the access area over the entire cargo runtime *τ* ^*c*^. Using the saturating value of *τ* ^*c*^ (for high *D*) in place of 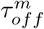 in the expression for *S*_*I*_ therefore gives an upper-bound estimate for the run length (upper dotted curve in Fig. 3(d)). We note that the top three curves for estimates that include the effects of reduced interference and increased availability of motors reflect the measured values of run length fairly well as a function of increasing surface diffusion constant across different conditions of ATP and motor number (see S16 Fig).

## Conclusion

Our results show that rigidly coupled motors experience a significantly broader range of forces when multiple motors are bound, reflecting the geometric constraints on the motor-cargo attachment points leading to antagonistic forces and therefore negative mechanical interference between motors. This interference is consistent with literature findings [28–30] and counteracts the expected load-sharing with increasing numbers of motors. We show that the increased fluidity of the lipid membrane which allows the attachment points to move and relax strain, decreases the negative interference between kinesins and permits load-sharing between motors to be more effective. The reduction of negative interference not only contributes almost equally to run length at low ATP (and/or high motor numbers) as increased motor availability but is, in fact, the dominant effect at saturating ATP and low motor numbers (see S16 Fig(a)) which is typically the normal physiological (and in vitro experimental) state. Direct measurements of this enhanced load-sharing may be possible using optical trap experiments with an applied load on rigid and fluid cargo. Tuning the load and motor number to a regime where enhanced load-sharing can significantly decrease motor off-rates may be able to resolve the effects in comparisons of rigid versus fluid cargo.

The effects of inter-motor interference in rigid cargo are also expected to be seen in measurements of speed. However, while our measurements of speeds from the simulations do show a decrease with increasing numbers of bound motors (S14 Fig), it is not as dramatic as reported in [41]. Consistent with [41], we find this decrease to be negated in the presence of fluidity but the effect is not statistically significant. One possibility is that the force-velocity relation we used (Eq. 11 in Materials and Methods) is not sensitive enough at low loads. Another more intriguing possibility is that the experimentally observed speed-up of lipid cargo [41, 42] may be due to a dispersion of speeds among motors leading to a bias for faster motors in the lead, and slower motors trailing and unbinding more often.

Since the reduction in interference manifests as a decrease in the off-rate of motors, it leads to increasing numbers of bound motors at higher membrane diffusion constants. Interestingly, we found that the on-rate was not constant, but increased with an increasing number of bound motors that effectively pulled the cargo surface closer to the microtubule. Monitoring the binding times of successive motors in an optical trap geometry could potentially allow for the experimental verification this effect. This interesting cooperative effect on the on-rate that we uncovered has a smaller effect, in general, on the run length than the reduced interference or increased availability of motors but is still significant (≈ 10%), especially at low ATP and/or high motor numbers (see S16 Fig). Thus, while the cooperative increase in on-rates works for both rigid and fluid cargo, the effect is enhanced when more motors are available due to fluidity. Finally, we showed that increased fluidity results in an expected increased recruitment of motors to the microtubule, thereby increasing the number of bound motors, with the effect being significant at very high motor density or low ATP. We found that the confluence of decreased interference, decreased off-rate, increased on-rate and increased motor recruitment can lead to positive feedback resulting in dramatic effects on the run length, under the right conditions. We show, for example, that run length is insensitive to cargo surface fluidity for moderate numbers of motors at saturating ATP but can increase by *several fold* for fluid membranes at low ATP conditions. Taken together, our work reconciles varying experimental results including the observed insensitivity of run length to cargo fluidity at moderate motor densities and high ATP concentrations [41, 42]. Our prediction of a significant enhancement of run length at low ATP is potentially physiologically important as an adaptive mechanism, *in vivo*, under stress or starvation conditions [47].

We developed a generalized version of the analytical expression for run length that accounts for cargo surface fluidity and expect it to be useful to explore a wide range of parameter spaces for different motor types and cargo geometries going beyond our simulations, which were done with a fixed cargo radius of 250 nm. Cargo geometry enters into the effective on-rate for motors which scales as *π*_*ad*_ = *π*_0_*S*_*a*_*/*(*S*_*I*_ + *S*_*a*_). The increase in access area, *S*_*a*_, as the cargo is pulled closer to the MT is determined by the curvature of the cargo surface and sets the magnitude of the cooperative increase in motor numbers. Even considering a fixed cargo height and high *D*, the on-rate still depends on the ratio of access area to the total surface area which decreases with increasing radius of curvature of the cargo. Thus, from our generalized version of Eq. 23, we expect an increased on-rate and higher run lengths for smaller cargo with a fixed number of motors (see S17 Fig), which is consistent with other numerical studies [36]. The smallest cargos, with sizes comparable to synaptic vesicles of about 100 nm [48], in fact, show an enormous increase in the run length over a range of fluidity that is consistent with the ranges for *in vivo* membranes in different contexts [38, 49]. For synaptic vesicles with fluid membranes therefore, transport over very large axonal distances, is a natural outcome of our model. If the surface density of motors is fixed, on the other hand, the quadratic increase in the number of available motors compensates for the reduction in on-rate resulting in larger run lengths for larger cargo (S18 Fig). In general, the changes in run length over physiological ranges of fluidity imply that surface fluidity could be used as a control parameter to regulate transport.

Our results on the transport properties of fluid cargo over a wide range of surface fluidity, ATP concentration, motor number and cargo size offer quantitative guidelines for the design of artificial cargo driven by kinesin teams. While all our results are specifically using parameters for kinesin motors, they can be readily generalized to other systems with different geometries of cargos with different motors coupled by a fluid surface. Thus our results could be of general value in designing artificial cargo utilizing teams of engineered versions of non-cooperative motors with fluid coupling to enhance load-sharing and processivity. More generally, coupling intrinsically non-cooperative mechanical force-generating elements via a fluid surface could be a generic mechanism to promote load-sharing that is exploited within the cell.

## Acknowledgments

We thank helpful discussions with David Quint and Jing Xu. This work was supported by the National Science Foundation (NSF-DMS-1616926) and NSF-CREST: Center for Cellular and Bio-molecular Machines at UC Merced (NSF-HRD-1547848). We also acknowledge partial support from the NSF Center for Engineering Mechanobiology grant CMMI-154857 and computing time on the Multi-Environment Computer for Exploration and Discovery (MERCED) cluster at UC Merced (NSF-ACI-1429783). NS acknowledges Graduate Student Opportunity Program Fellowship from the University of California, Merced.

## Materials and methods

We developed a Brownian dynamics model to address the question of how cargo surface fluidity influences transport by teams of kinesin motors. Early models of multi-motor cargo transport used a mean-field approach where all motors were considered to share the load equally [50]. Later stochastic models that assume unequal load sharing [30, 40, 43, 51–54] were found to describe experimental observations better [43, 55, 56]. The next improvement to the model was to explicitly consider that motors are bound to a three dimensional spherical cargo surface, that was typically considered rigid [57, 58]. Only a couple of recent models have attempted to consider the lipid membrane on the cargo surface [36, 42].

Our model is a stochastic model that considers unequal load sharing with explicit implementation of motor diffusion on the cargo surface. In our model, we consider a spherical cargo of radius, *R* = 250 nm which is decorated with a given number of molecular motors (*N*) on the surface. *N* doesn’t change during the cargo run, meaning there is no binding and unbinding of motors between the cargo surface and the solution. Each of these *N* motors is initially assigned a random, uniformly distributed anchor position on the cargo surface. Molecular motors on rigid cargo are fixed on the cargo surface. So a given initial configuration of motors on the rigid cargo surface persists throughout the cargo run whereas motors on lipid cargo diffuse on the surface.

In this study we considered that all *N* motors are kinesin motors which have a rest length of *L*_*mot*_ = 57 nm [1]. We assume that an unbound kinesin motor binds to the microtubule with constant rate, *π*_0_ = 5 s^−1^ [40], if some part of the microtubule is within *L*_*mot*_ distance from the anchor position of that unbound motor. In other words, unbound motors bind with a specific rate to the microtubule if they can access the microtubule.

A microtubule bound motor is characterized by two position vectors, anchor point 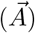 on the cargo surface and head 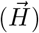 on the microtubule. A microtubule bound motor is assumed to exert a spring-like force when the length of motor exceeds the motor’s rest length (*L*_*mot*_) with a force constant, *k*_*mot*_ = 0.32 pN/nm [59, 60]. The force exerted by a motor on the cargo is then given by,

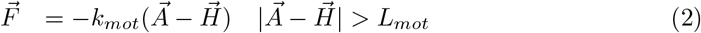

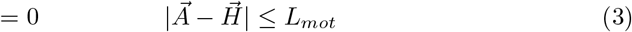

The translation velocity of the center of mass of cargo is given by the (overdamped) Langevin equation

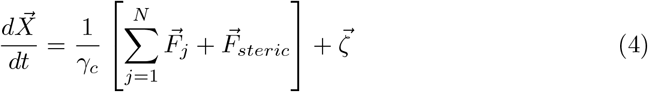

Here, 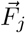 is the force exerted on cargo by the j^*th*^ motor. 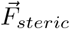 is the spring-like steric force on the cargo from the microtubule, represented with a high force constant 10*k*_*mot*_. We note that if the vesicle is deformable the steric spring constant could be significantly smaller. This force is present only if the cargo-microtubule distance is less than the sum of the cargo and microtubule radii. 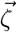 is the random force on the cargo due to collisions with the intracellular medium. We assume a normally distributed noise with zero mean, 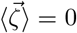 and the fluctuation-dissipation relation, ⟨ *ζ*_*μ*_*(t)ζ*_*ν*_*(t′)*⟩ *= 2γ*_*c*_*k*_*B*_*Tδ*_*μν*_*δ(t*−*t*′). *γ*_*c*_ is the friction co-efficient for the cargo given by *γ*_*c*_ = 18*πη*_*v*_*R*, where *η*_*v*_ is the co-efficient of viscosity of cytoplasm experienced by cargo. We approximated *η*_*v*_ to be equal to the co-efficient of viscosity of water, *η*_*v*_ = 10^−3^ Pa.s. We integrate Eq. 4 using the Euler-Maruyama scheme

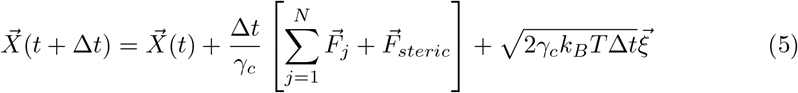

where 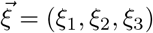 and *ξ*_1_, *ξ*_2_, *ξ*_3_ are drawn from normal distribution with zero mean and unit variance.

At each time step we also update the anchor positions of each motor on the lipid cargo surface using a similar Brownian dynamics formalism given by

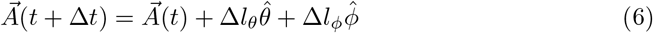

where Δ*l*_*θ*_ and Δ*l*_*ϕ*_ are the small displacements along 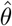 and 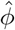 directions in the plane tangential to cargo surface at 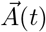. Δ*l*_*θ*_ and Δ*l*_*ϕ*_ are given by

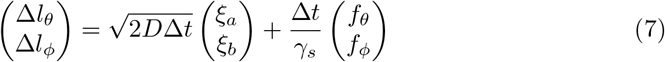

where *ξ*_*a*_ and *ξ*_*b*_ are random variables obtained from normal distribution with zero mean and unit variance. *D* is the diffusion constant for motor diffusion on cargo surface and *γ*_*s*_ is the friction coefficient given by *γ*_*s*_ = *k*_*B*_*T/D*. (*f*_*θ*_, *f*_*ϕ*_) are the components of motor forces along 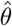 and 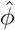 respectively which can be obtained from the motor force in Cartesian co-ordinates, 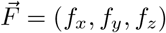 as follows

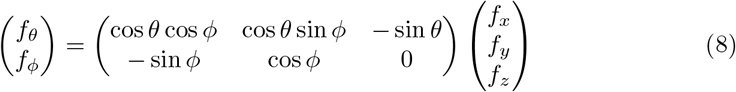

At every time step, each bound kinesin motor hydrolyses an ATP molecule with certain probability and attempts to move forward on the microtubule. This stepping probability is a function of the motor force and also the ATP concentration. We have adopted the following relation for the stepping probability [1]

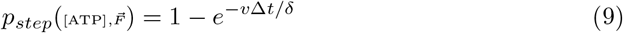

where *δ* is the step size, the distance moved by motor after hydrolyzing one ATP molecule (*δ* = 8 nm [13, 61, 62]). *v* is the velocity of kinesin motor which is a function of the ATP concentration and motor force and is taken to be

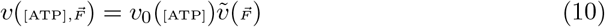

where *v*_0_(_[ATP]_) is the velocity of motor under no-load condition at a given ATP concentration. The ATP dependence of *v*_0_ is described by the Michaelis-Menten equation

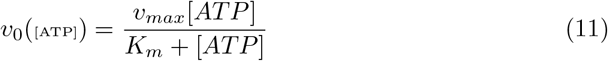

We considered the no-load velocity at saturated ATP concentration to be *v*_*max*_ = 800 nm/s [1, 2] and *K*_*m*_ = 44 *μ*M [3].

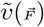 gives the force dependence of the velocity. In the hindering direction, we assume [1, 43]

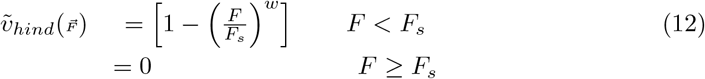

F is the magnitude of motor force, 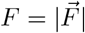. *F*_*s*_ is the stall force, the value of force beyond which kinesin motor stops walking. We considered *F*_*s*_ = 7 pN [2, 64, 65] and *w* = 2 [55]. In the assistive direction, velocity is assumed to be independent of force magnitude, 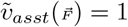 [1, 2].

Experimentally it is found that a kinesin motor is more likely to detach from the microtubule when one head is detached from the microtubule while trying to take a step than when both the heads are bound to the microtubule [3–5]. In our model we assume that a motor can detach only when it tries to take a step. At every time step we first check whether a bound motor tries to make a step with probability 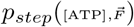 using Eq. 9. If it tries to take a step, we check whether it detaches from the microtubule before completing the step using a microscopic off-rate whose value is calibrated based on the experimentally observed off-rate as a function of force, *F*, at saturating ATP concentrations.

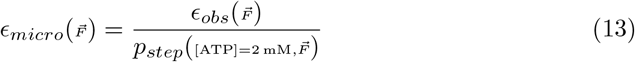

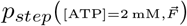 is the probability to step forward in time step Δ*t* at high ATP concentration of 2 mM.

For hindering forces, we used the following relationship between observed off-rate and magnitude of motor force *F* developed in [15] based on Kramer’s theory [68] and used in several studies [1, 23, 43]

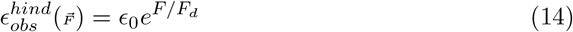

*ϵ*_0_ is the off-rate under no load condition. We used *ϵ*_0_ = 0.79 *s*^−1^ [1, 2]. *F*_*d*_ is the detachment force. We approximated *F*_*d*_ to be equal to the stall force *F*_*s*_. For assistive forces, the relationship between observed off-rate and magnitude of force is taken to be [1, 2]

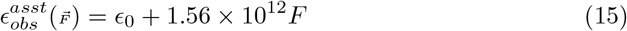

## Supporting information

**S1 Fig.**
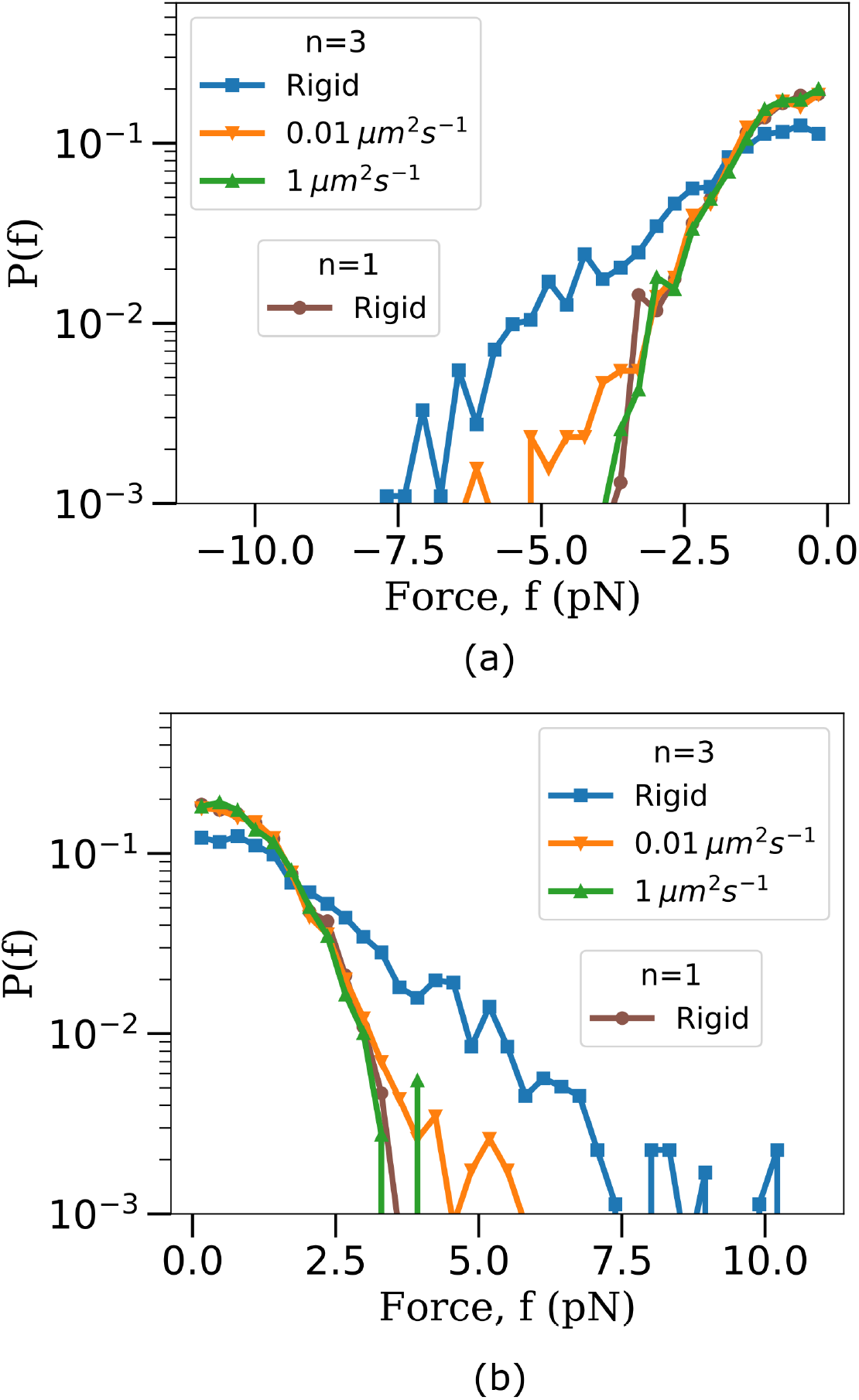
Distribution of forces separately in (a) hindering direction and (b) assistive direction. n represents the number of motors bound to cargo. The procedure followed in obtaining the force values is the same as explained in Fig. 2 of the main paper. These distributions show that, in a rigid cargo, there is negative interference between multiple motors both in assistive and hindering directions and the lipid membrane reduces this negative interference.

**S2 Fig.**
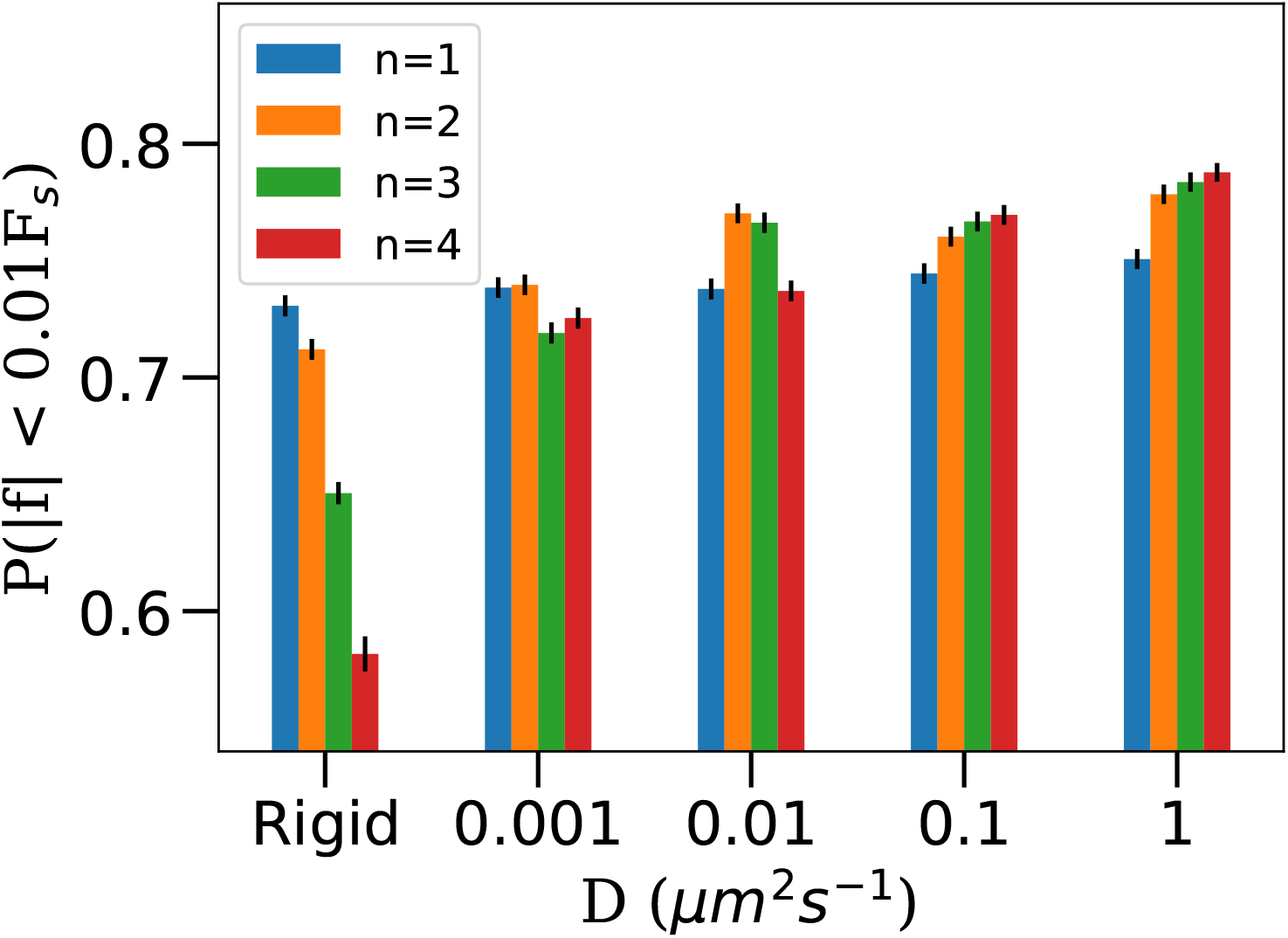
The fraction of the force distribution with a magnitude of force *f* greater than or equal to 0.01*F*_*s*_, as a function of the diffusion constant for the different number of bound motors n. Error bars represent the standard error of the mean obtained by the bootstrap method. *F*_*s*_ is the stall force of kinesin. The fraction increases as a function of n for rigid cargo but decreases for lipid cargo. Required force distributions as a function of *n* and *D* were obtained using the same procedure as explained in Fig. 2.

**S3 Fig.**
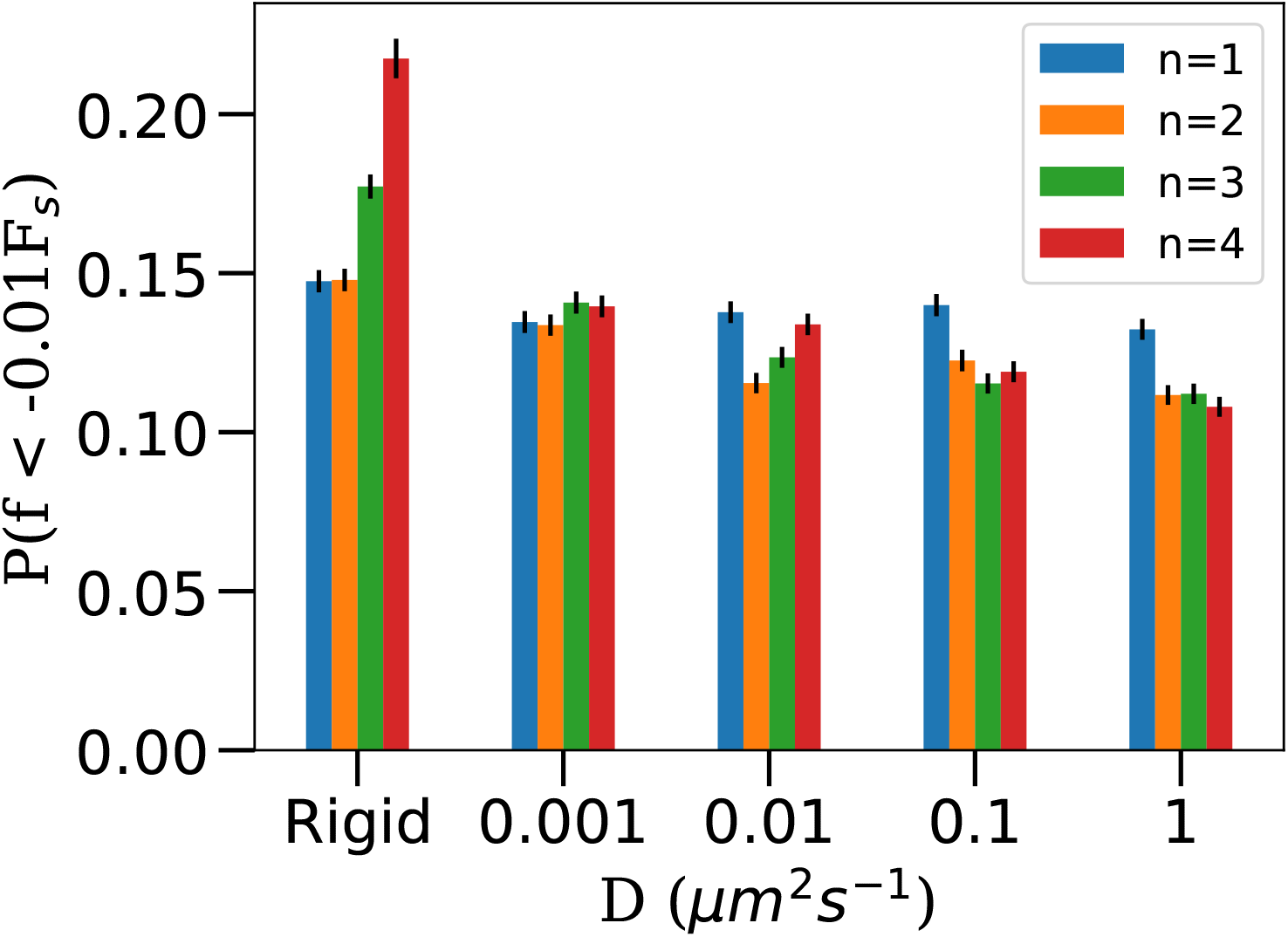
Fraction of force distribution with force magnitude greater than 0.01*F*_*s*_ in the hindering direction. Required force distributions as a function of *n* and *D* were obtained using the same procedure as explained in Fig. 2.

**S4 Fig.**
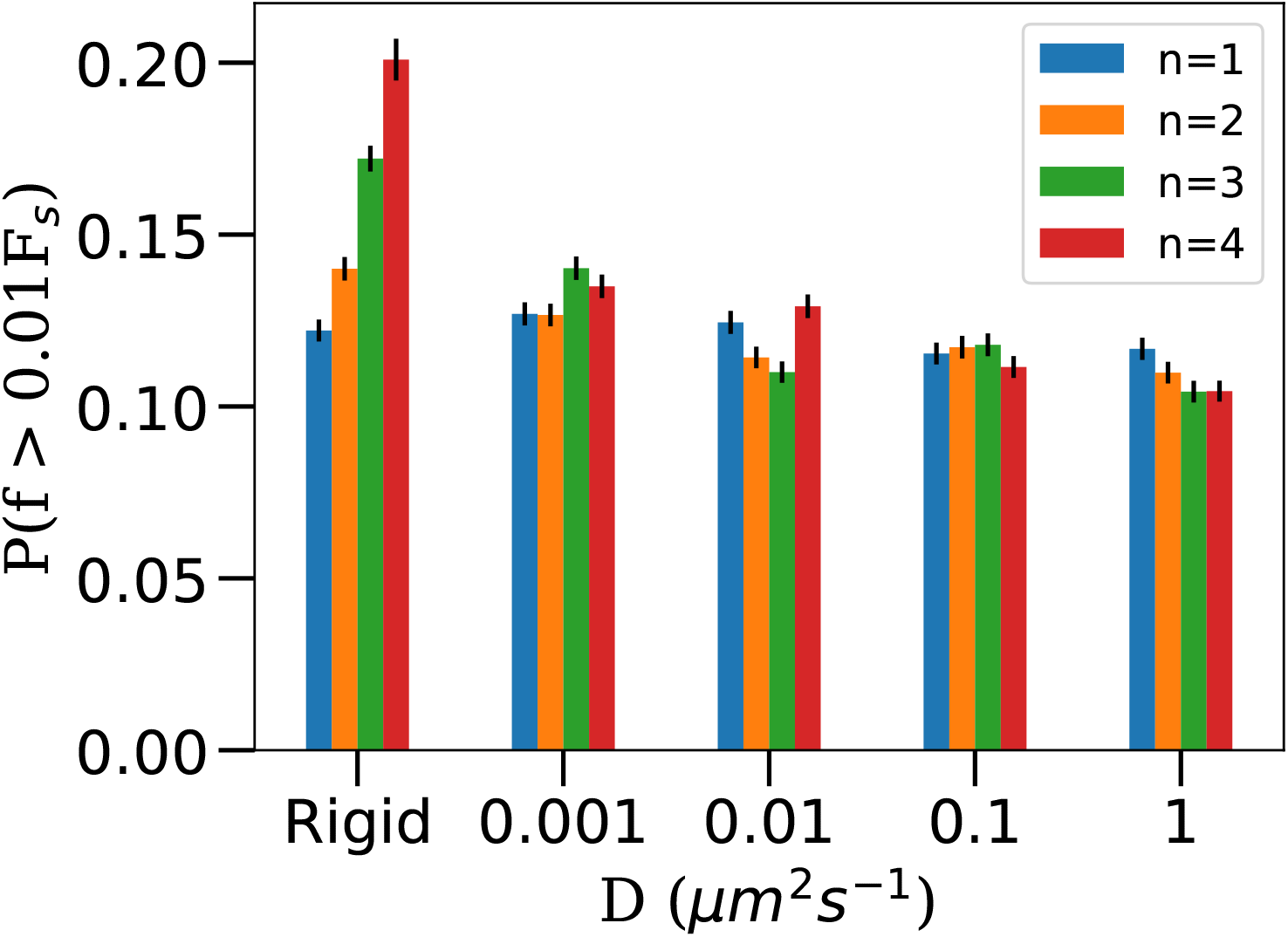
Fraction of force distribution with force magnitude greater than 0.01*F*_*s*_ in the assistive direction. Required force distributions as a function of *n* and *D* were obtained using the same procedure as explained in Fig. 2.

**S5 Fig.**
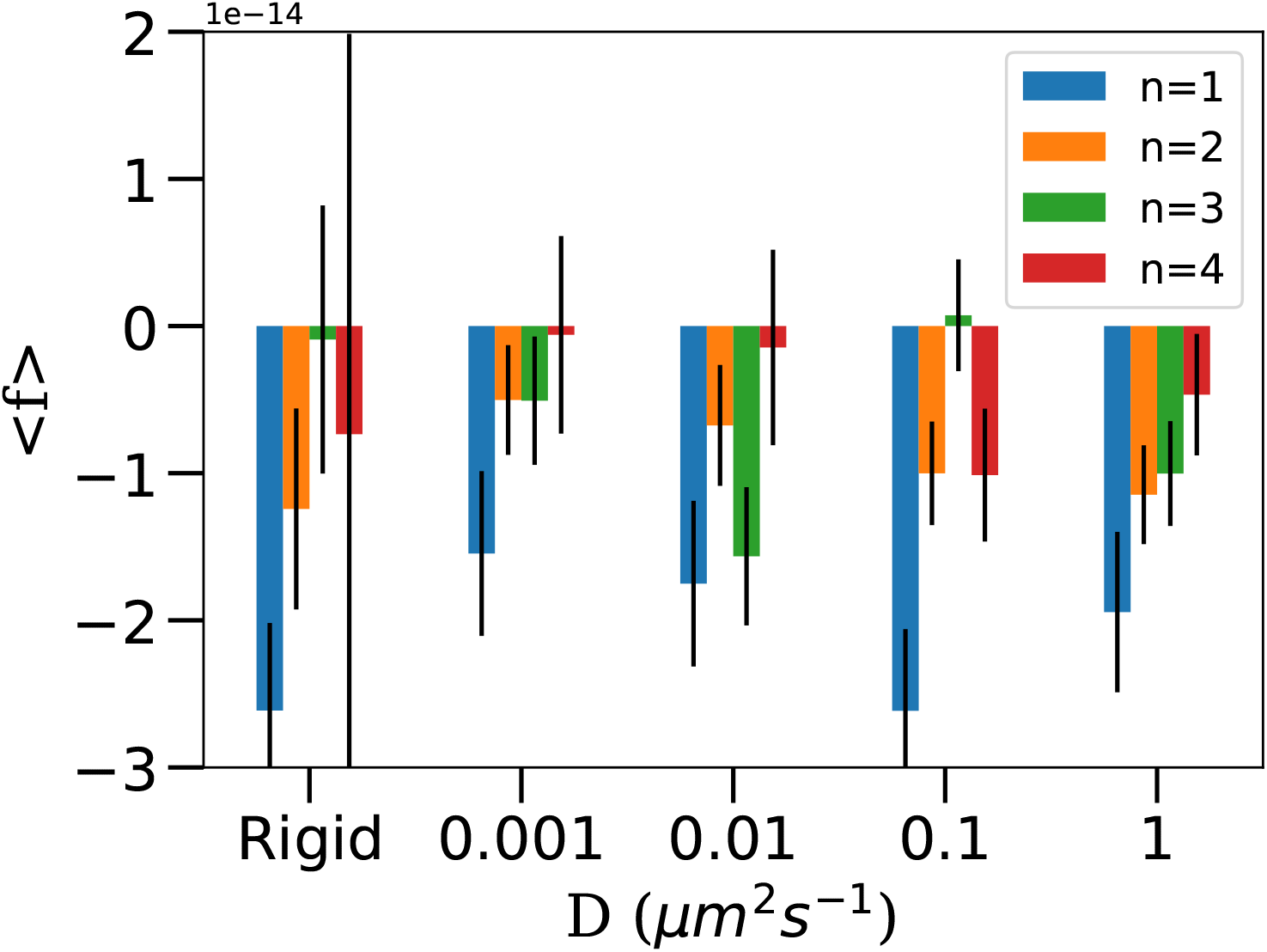
Mean value motor force (in Newtons) experienced by a bound motor as a function of the number of bound motors *n* and diffusion constant *D*. Mean force is in hindering direction, as one would expect because there is an active motion of motor in that direction. The magnitude of mean force decreases with an increase in the number of bound motors due to load sharing. Force distribution data is the same as Fig. 2 of the main paper.

**S6 Fig.**
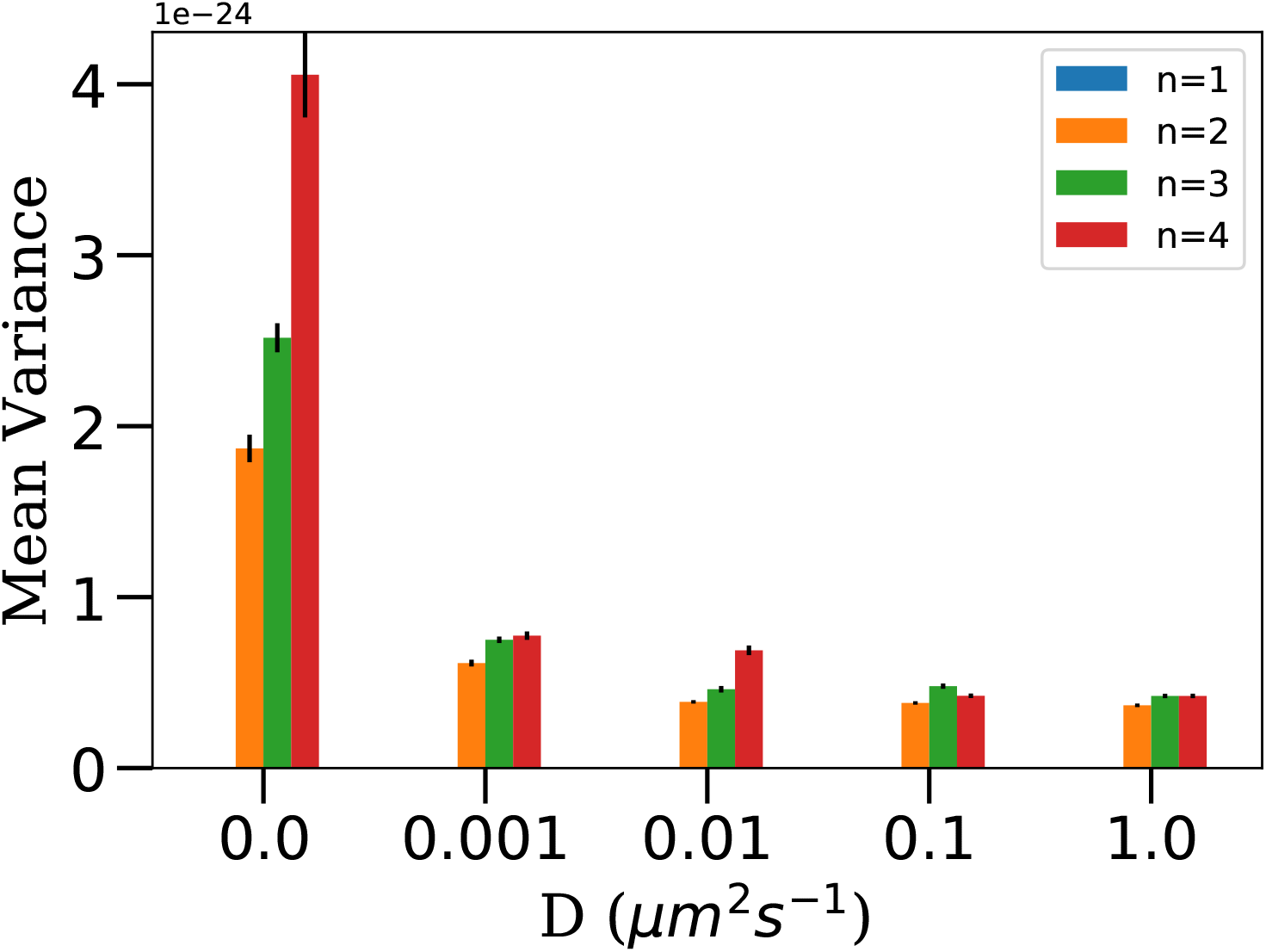
The mean variance of forces (in units of *N* ^2^) among bound motors. Calculated as 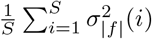 where the summation is over all the sample time points where the number of bound motors is *n*. 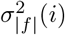 is the variance in the magnitude of force experienced by the n bound motors at *i*^*th*^ data sample. *S* = 10000 except for (i) *D* = 0 *n* = 3, *S* = 7360 (ii) *D* = 0 *n* = 4, *S* = 1083 (iii) *D* = 0.001 *n* = 4, *S* = 3754 (iv) *D* = 0.01 *n* = 4, *S* = 6396 (v) *D* = 0.01 *n* = 4, *S* = 3508 (vi) *D* = 0.1 *n* = 4, *S* = 4883. Motor force data is from the same simulation described in Fig. 2 of main paper (N = 16 [ATP] = 2 mM).

**S7 Fig.**
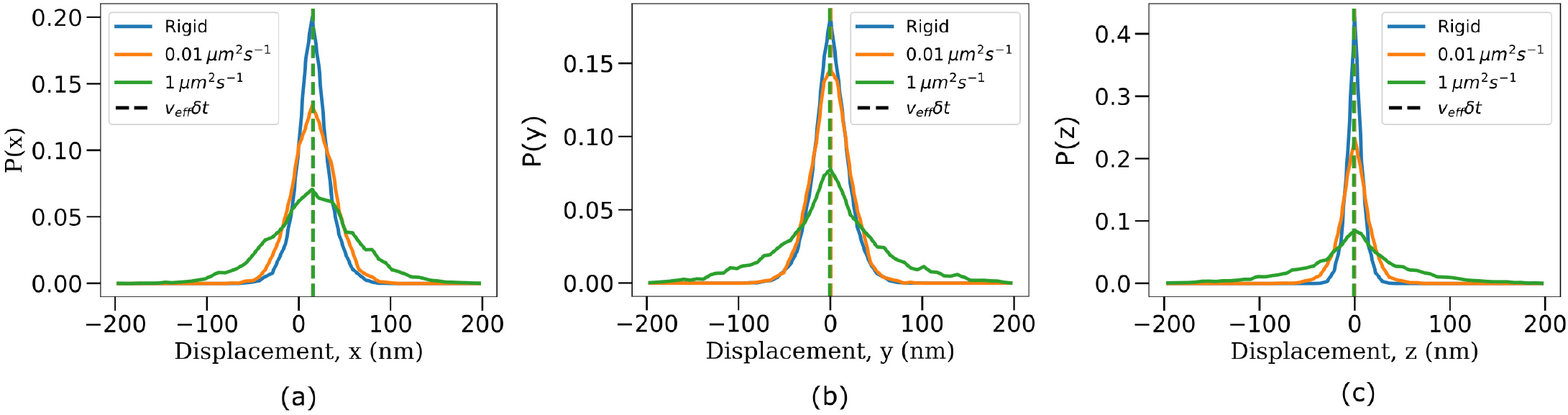
Distribution of cargo displacements (position fluctuations) measured in 0.01 s time intervals along (a) x-axis (b) y-axis and the (c) z-axis. In general, fluctuations increase with the increase in the fluidity of the cargo surface. Vertical lines indicate the mean values of fluctuations, interpreted as *v*_*eff*_ *δt* where *v*_*eff*_ is the effective velocity of cargo and *δt* is the time interval used for measuring fluctuations (0.01 s). Fluctuations along the x-axis have a positive mean (*v*_*eff*_ *δt* ≈ 8*nm*) because of the active motion due to molecular motors. Fluctuations along the y-axis and z-axis have a 0 mean. Fluctuations along the z-axis are much narrower than fluctuations along the y-axis and x-axis because of the steric force due to the microtubule. Data were obtained from the transport of cargoes with a total of *N* = 16 motors at [ATP] = 2 mM. 200 cargo runs were considered for each diffusion constant.

**S8 Fig.**
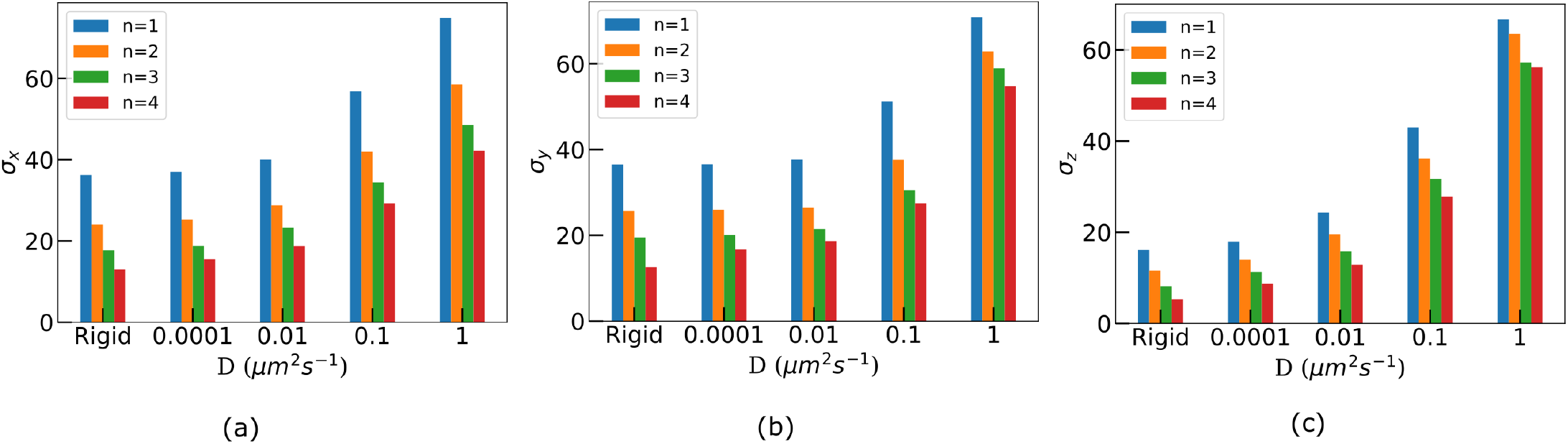
Standard deviation (nm) of the position fluctuations as a function of diffusivity and the number of bound motors. Data was obtained from the simulation of the transport of cargoes with a total of *N* = 16 motors at [ATP] = 2 mM. 200 cargo runs were considered for each diffusion constant.

**S9 Fig.**
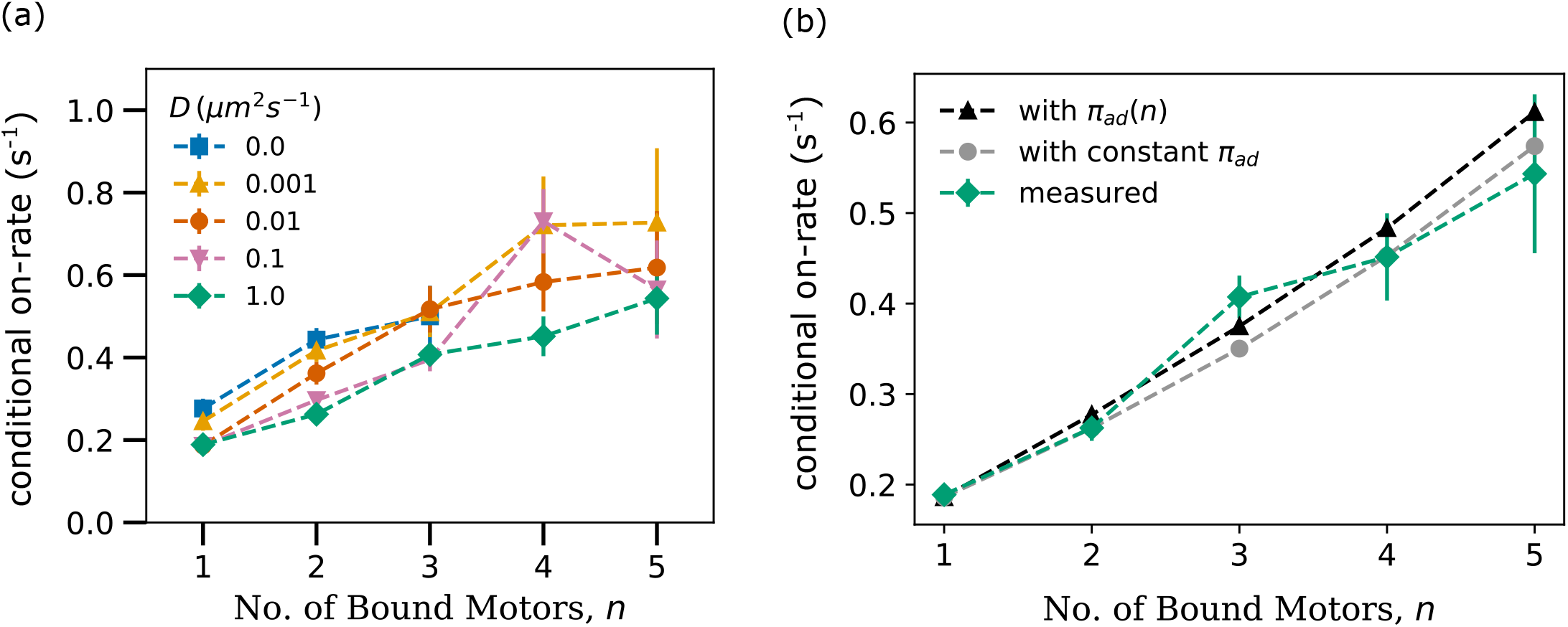
**(a) Conditional on-rate measured from simulation trajectories.** We simulated 200 cargo runs for N = 16 [ATP] = 2 mM case and recorded the value of the number of bound motors at a sampling rate of 100 s^−1^. With this data, we identified all the *n* bound motors to *n* + 1 bound motor transitions, calculated the mean time for such transitions, and then the mean rate (1/meantime). We then divided this rate by the number of unbound motors *N* − *n* to obtain the conditional on-rate per motor. We then repeated the analysis process for a different number of bound motors *n* and diffusion constants *D*. **(b) Comparing the measured conditional on-rate with analytical expressions**. Black triangles with dashed lines provide conditional on-rate where we consider the increase in on-rate of a single motor with an increase in the number of bound motors. Grey circles with a dashed line provide conditional on rate assuming a constant single motor on-rate. It can be inferred that if we measure this quantity in experiments one can expect to see only a slight increase as the cargo comes closer to the microtubule. This is because as the number of bound motors increases, the effective detachment rate of the n-bound state increases, hence the gating time decreases. Our analysis shows that the decrease in gating time contributes more to the increase in the conditional on-rate than the increase in the single motor on-rate. See S2 Appendix for more details.

**S10 Fig.**
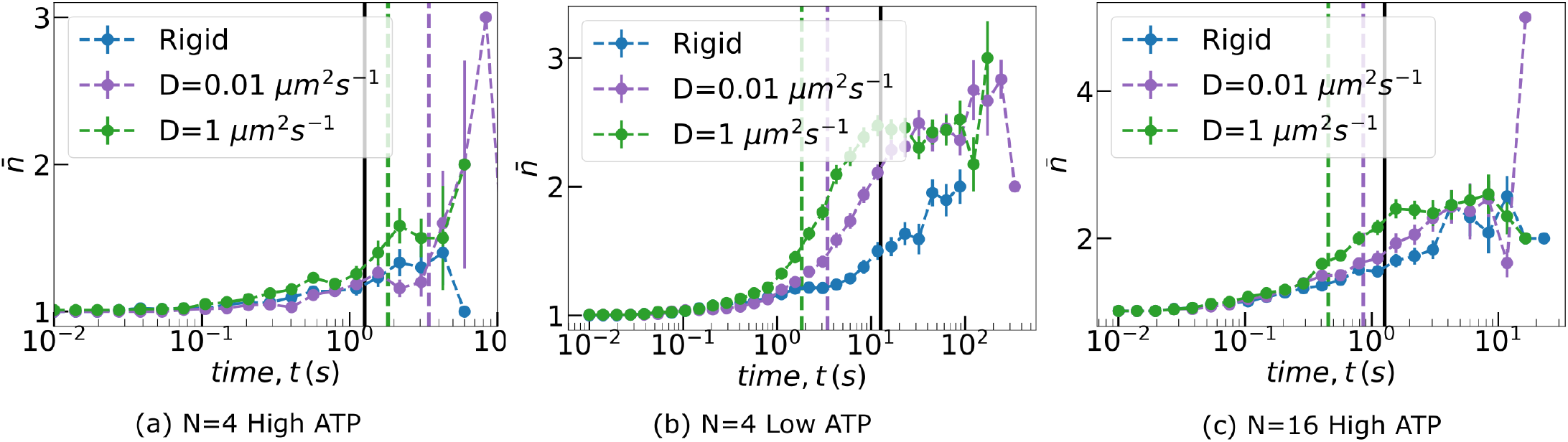
Ensemble average (± SEM) of the number of bound motors for three different cargo-motor systems. Vertical lines indicate the estimated time-scales, mean time for a new motor to bind - *τ*_*bind*_ (dashed lines) and mean unbinding time of a kinesin motor - *τ*_*off*_ (solid black line). N is the total number of motors on cargo, High [ATP] = 2 mM, Low [ATP] = 4.9 *μ*M. The averaging was performed over 200 cargo runs each. Please refer to Table I in main text for more information on *τ*_*bind*_ and *τ*_*off*_.

**S11 Fig.**
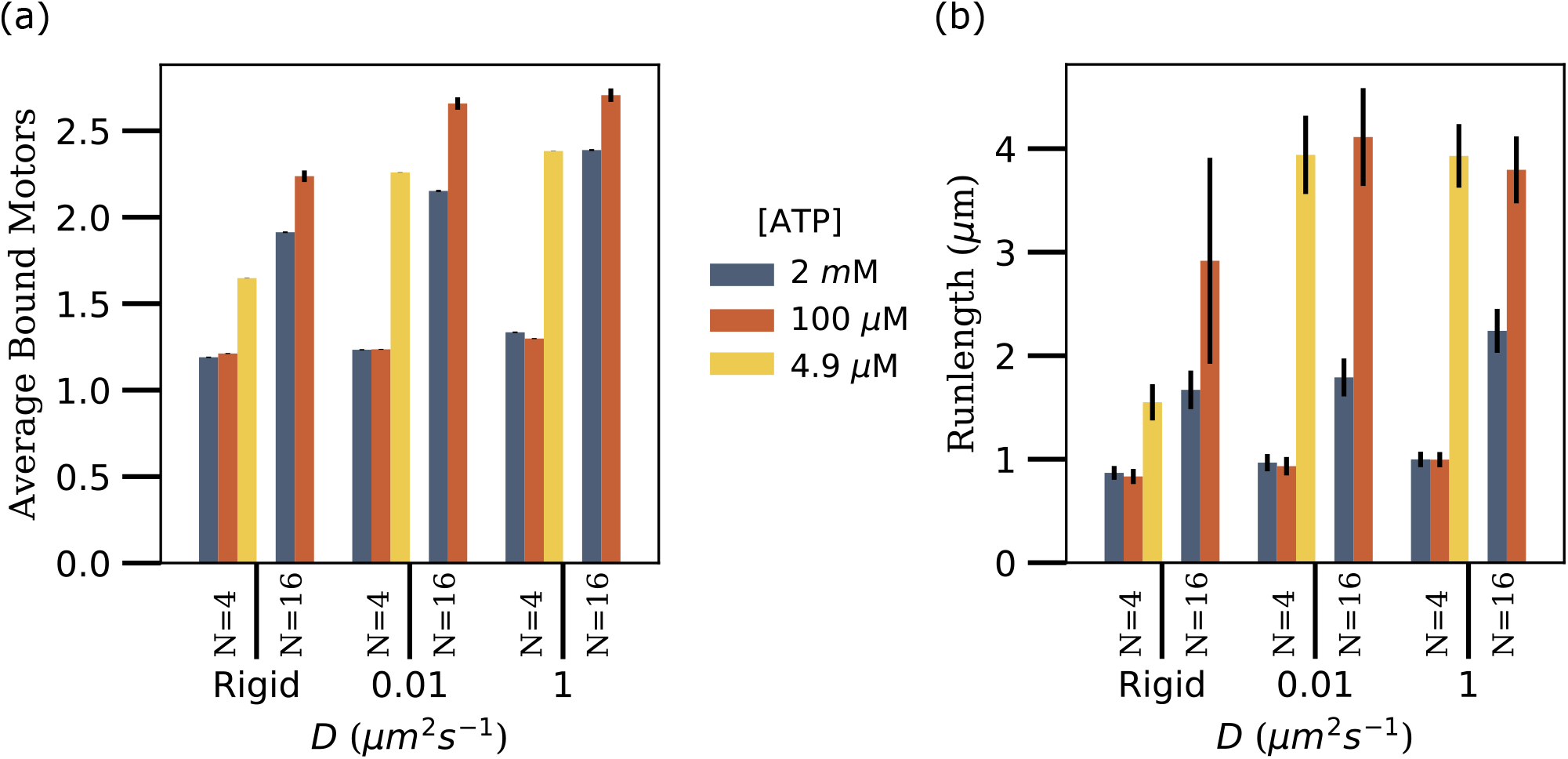
(a) Average number bound motors, (b) Runlength as a function of diffusivity (*D*) for two different numbers of motors on the cargo (*N*) and three different ATP concentrations. 200 cargo runs were considered for each parameter set. Error bars represent the standard error of the mean (SEM). The average number of bound motors in (a) was calculated as the mean of the number of bound motors (*n*) in all time samples (with data sampling rate = 100 s^−1^) of these 200 cargo runs. Thus the sample size was large and hence the error bars obtained as the standard error of the mean is very small. Runlength figure is same as the Fig. 3(c) in the main text, added here just to make it easy to compare.

**S12 Fig.**
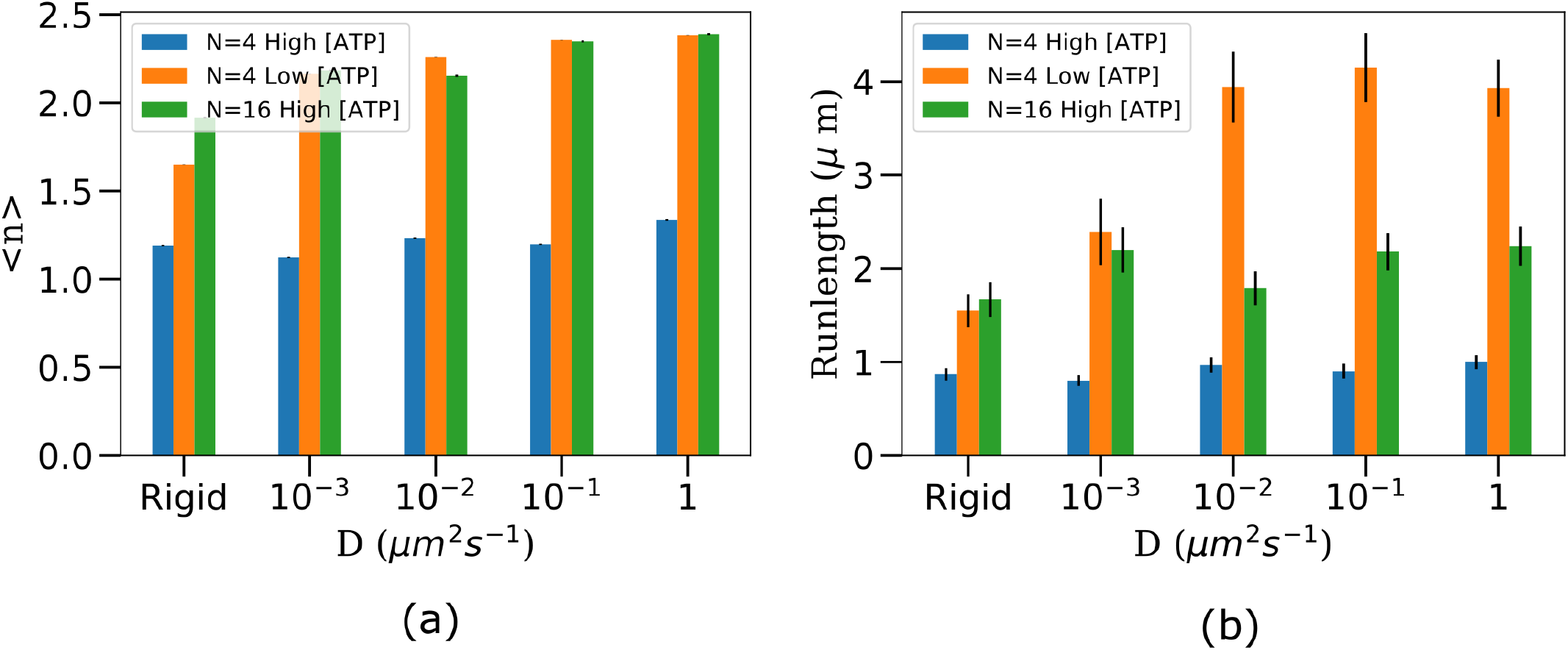
(a) Average number of motors (± SEM) (b) Runlength (± SEM) for more values of diffusion constants. Averaging was performed over 200 cargo runs in each case.

**S13 Fig.**
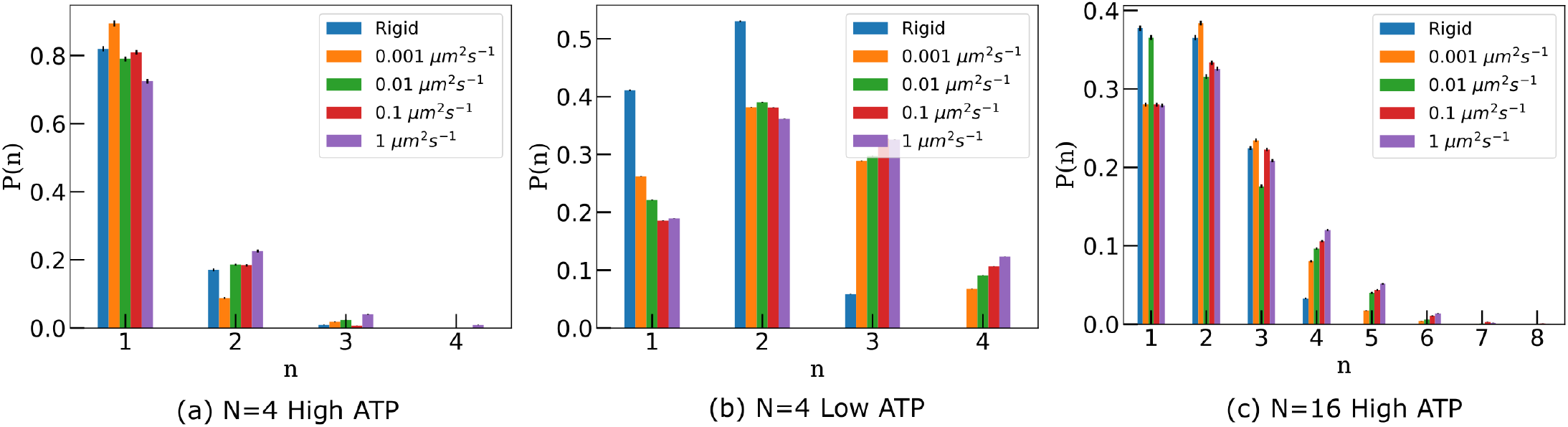
Probability distribution of the number of bound motors in three different cargo-motor systems. High [ATP] = 2 mM, Low [ATP] = 4.9 *μ*M. Data was obtained from 200 cargo runs for each case.

**S14 Fig.**
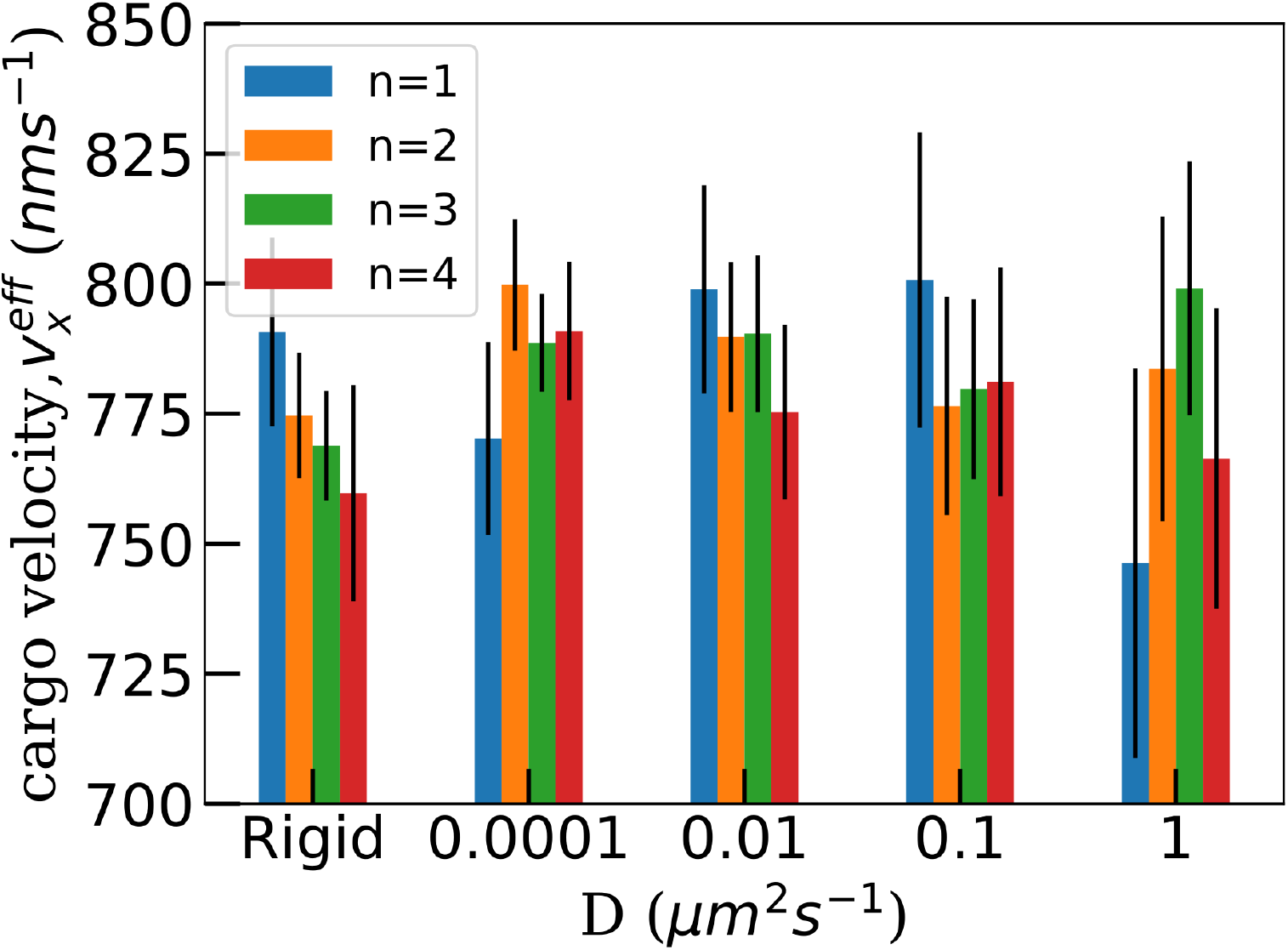
Velocity of cargo along x-axis, 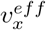 as a function of the number of bound motors and diffusion constant. Data was obtained from the simulations of the transport of cargoes with a total of *N* = 16 motors at [ATP] = 2 mM. 200 cargo runs were considered each for each diffusion constant.

**S15 Fig.**
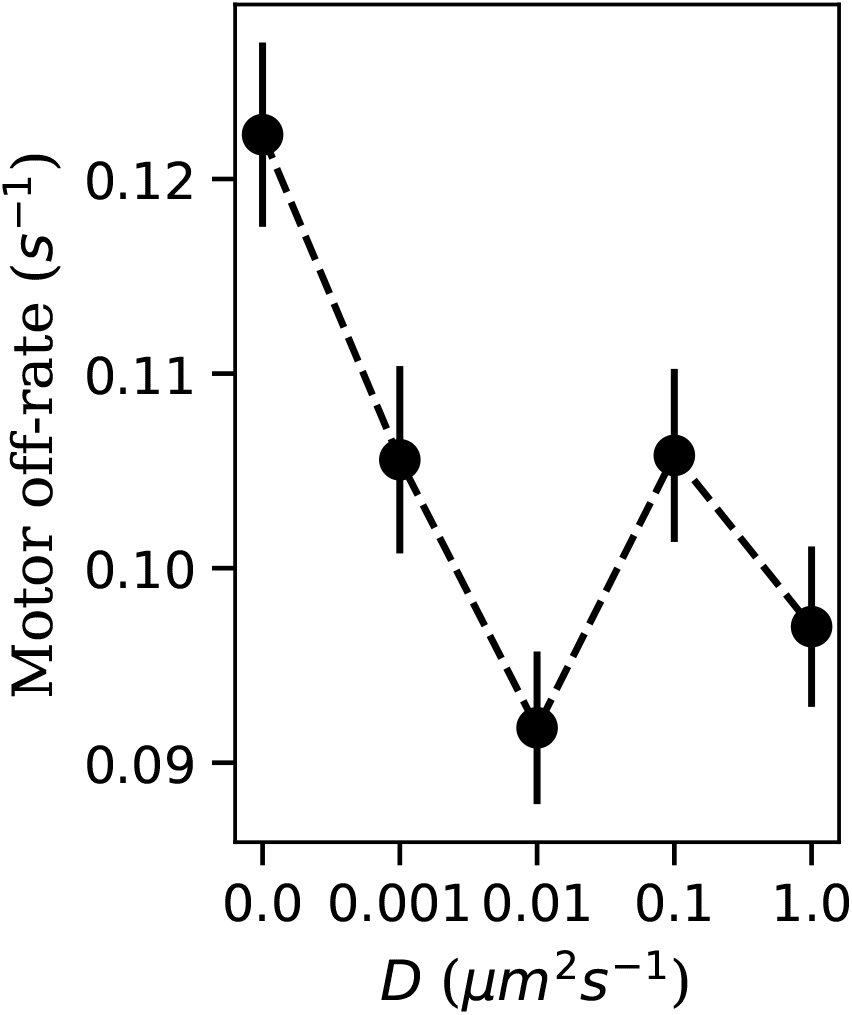
Average value of motor off-rate as a function of fluidity of cargo surface for N = 4 [ATP] = 4.9 *μM*. (Sample size = 530 from 200 cargo runs)

**S16 Fig.**
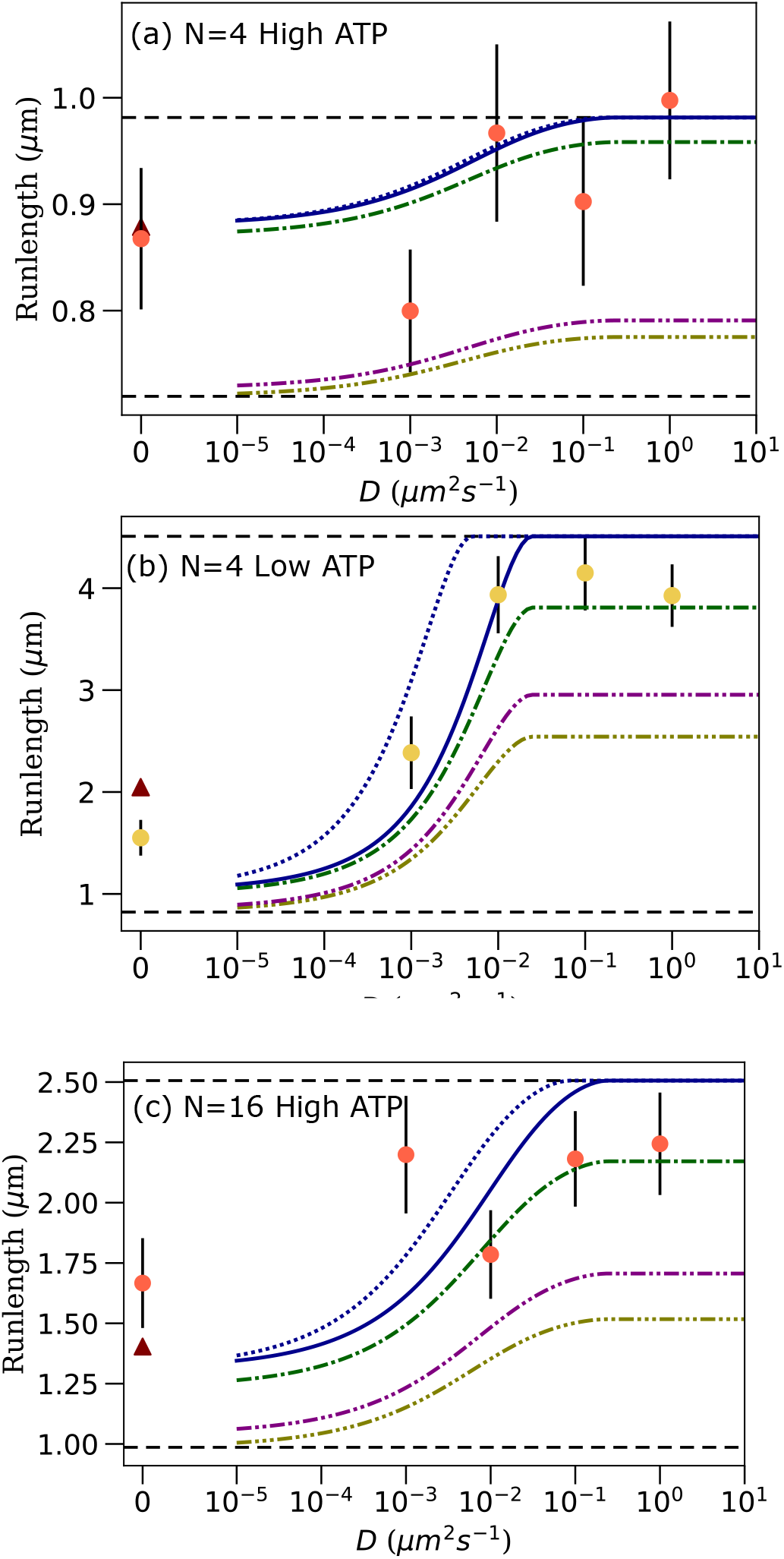
Comparison between run lengths from our simulations (circles) and analytical estimates (maroon triangle, solid, dashed and dash-dotted lines) as described in the main text.

**S17 Fig.**
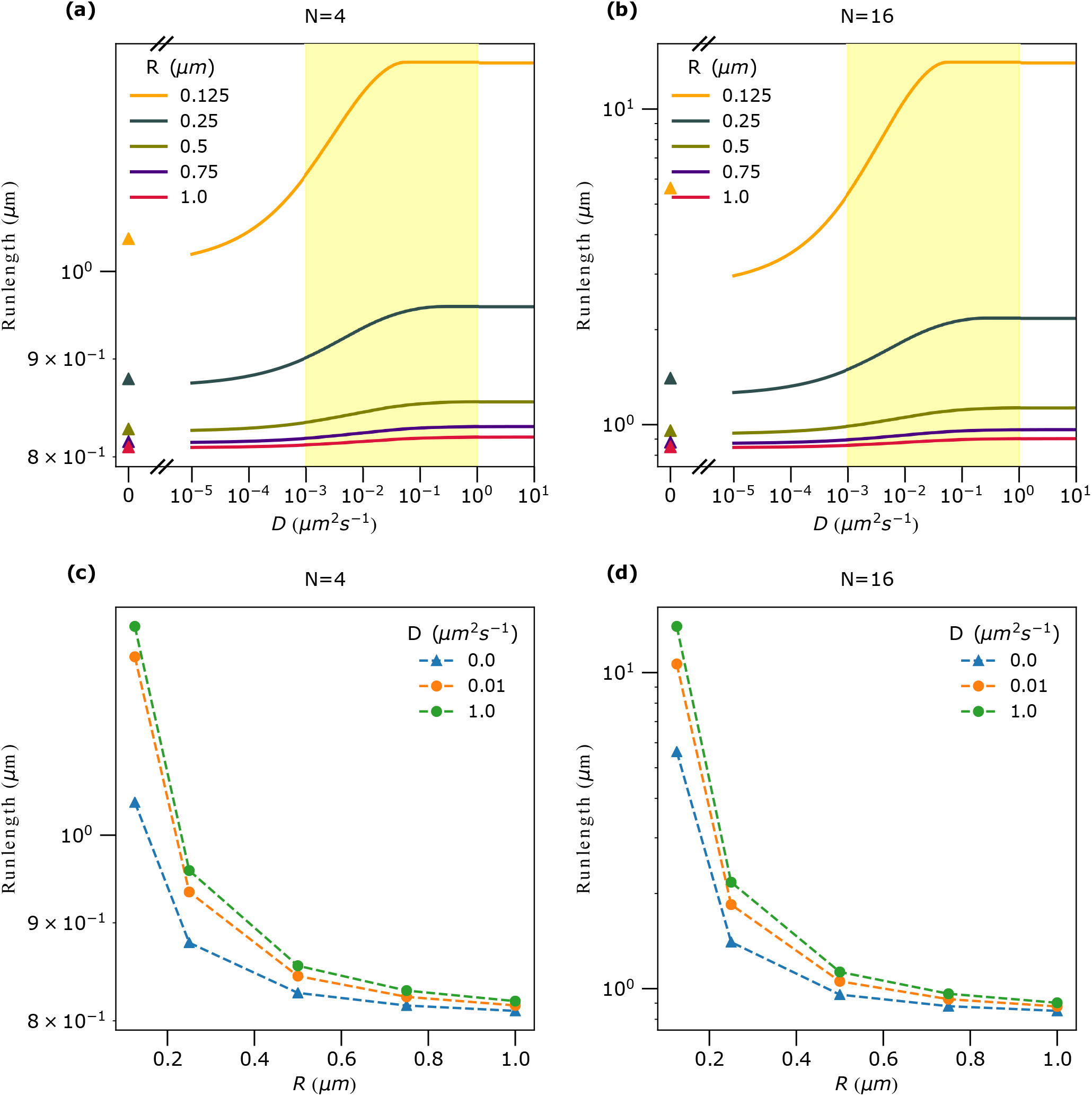
Analytically estimated runlengths for different cargo radii for a fixed number of motors on cargo. The general method for calculating runlength analytically is described in the main text. We have numerically computed access area, *S*_*a*_, considering the typical distance between cargo surface and MT for 1 motor bound case. 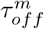 for computing influx area, 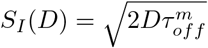, was taken to be 1 s which is the typical motor unbinding time at saturating ATP concentration. Yellow vertical bands in (a) and (b) correspond to the range of physiologically relevant diffusion constants of motors.

**S18 Fig.**
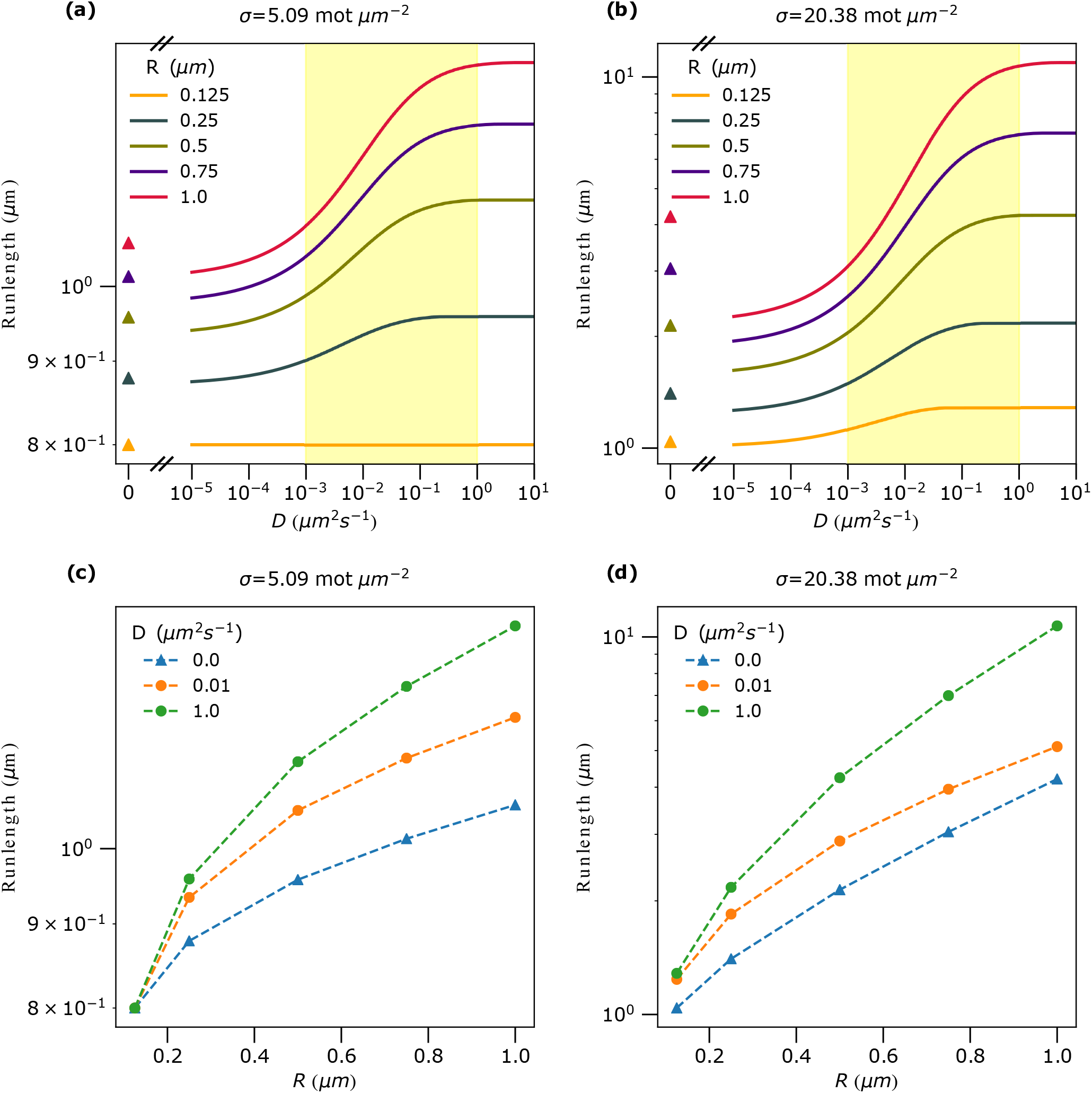
Analytically estimated runlengths as a function of cargo radius (R) for a fixed surface motor density (*σ*). The general method for calculating runlength analytically is described in the main text. We have numerically computed access area, *S*_*a*_, considering the typical distance between cargo surface and MT for 1 motor bound case. 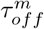 for computing influx area, 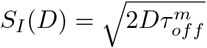, was taken to be 1 s which is the typical motor unbinding time at saturating ATP concentration. Yellow vertical bands in (a) and (b) correspond to the range of physiologically relevant diffusion constants of motors.

**S19 Fig.**
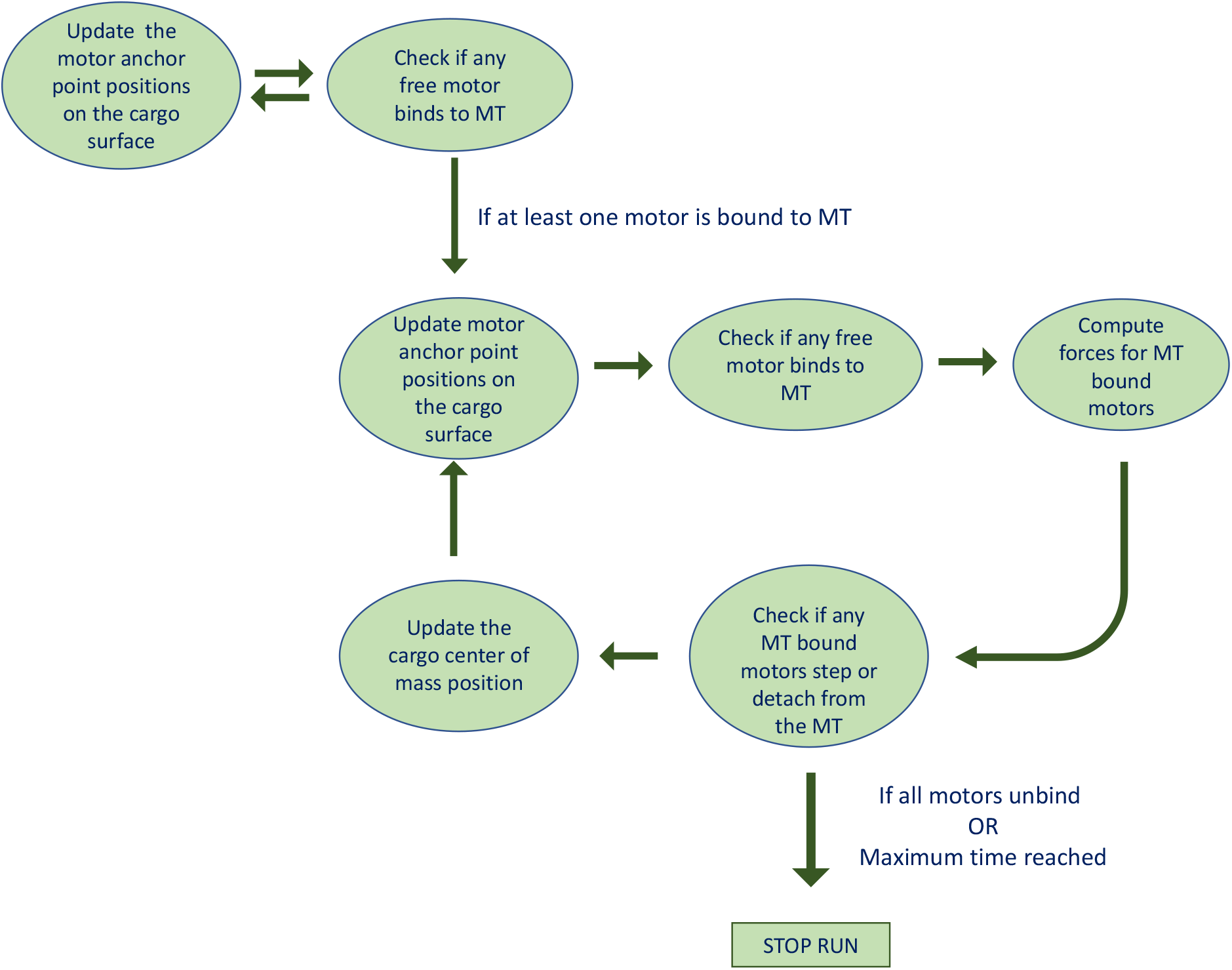
The flow chart of the simulation. The cargo is held near the microtubule until at least one motor is bound. Once a motor is bound, we loop over the series of steps shown in the middle until either all motors are unbound or maximum time is reached. At regular time intervals, we record relevant data like the cargo center of mass, anchor positions of motors on cargo surface, head positions of bound motors on the microtubule.

**S1 Video. Visualization of our cargo transport model**. The light brown colored sphere represents the cargo. Small red spheres are unbound motors. Unbound motors diffuse on the cargo surface and bind to the microtubule with a rate of 5 s^−1^ when they come close to the microtubule. Blue sticks represent bound motors that walk along the microtubule with a rate dependent on the force that they experience. Bound motors unbind with a force-dependent rate. The microtubule is represented as the dark-green horizontal line. Data for this movie was obtained from the simulation of a cargo with 16 motors. The data sampling rate was 10^5^ s^−1^. We used Mayavi [69] for visualization.

**S2 Video. Cargo dynamics viewed at a shorter time and length scale**. The data for this movie is the same as S1 Video.

## S1 Appendix

### Notes on the assumptions in the computational model

#### We neglect rotational diffusion of cargo

We expect the rotational diffusion to be minimum for a rigid cargo, increase with an increase in cargo surface fluidity and approach the value for free bead in solution at very high cargo surface fluidity. Nonetheless, one can show that such rotational diffusion of cargo only renormalizes the diffusion constant of motors on the surface (at most by a factor of 2) but doesn’t induce any qualitative change in the motor availability at the access region for any cargo surface fluidity.

Next we note that a rigid cargo can still rotate about the motor stalk axis [1], although we expect it to be possible only when a single motor is bound. The typical value of angular diffusion constant for this process was measured [1] to be *D*_0.5 *μ*m_ = 0.5 × 10^−2^ rad^2^*s*^−1^ for a bead of half-micron radius. Since angular diffusion constant is inversely proportional to *R*^3^, *D*_0.25 *μm*_ = 4 × 10^−2^ rad^2^s^−1^. This diffusivity is low to have any impact on the transport process. For example, the time required for a cargo to rotate by 90^*o*^ with this diffusion constant is 30.8 s which is greater than the lifetime of a single kinesin motor at even the lowest ATP concentration considered in our study. (lifetime of kinesin is about 1 s at [ATP] = 2 mM and about 10 s at [ATP] = 4.9 *μ*M). Hence we neglected the rotational diffusion of rigid cargoes also.

#### We neglect rotation of cargo due to torque from motor forces

When a motor pulls a cargo, often there is some component of force tangential to the cargo surface. This tangential component of force results in a torque on the cargo. We computed the typical magnitude of the angular velocity of the cargo by such torque and concluded that this angular velocity is too low to have any impact on the transport process.

Assume that a microtubule bound motor exerts a torque 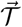 on the cargo. Angular velocity of cargo due to this torque is

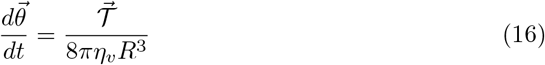

Typical value of tangential component of motor force is *f*_*tan*_ = 1 pN. So the typical magnitude of the torque on the cargo due to this force is 𝒯 = *Rf*_*tan*_ = 2.5 × 10^−19^ Nm. Substituting this in Eq. 16 we get the typical magnitude of angular velocity due to torque from motor forces to be equal to 3.98 × 10^−11^ rad s^−1^. Thus the rotation of the cargo due to motor forces in 1 s is 3.98 × 10^−11^ rad which corresponds to an arc length much lesser than 1 nm. So we can neglect cargo rotation due to motor forces in our computational model.

#### We neglect the deformation of vesicle due to motor forces

Application of a point force on the vesicle (the cargo) surface leads to shape deformation which could develop as narrow membrane tubes (tethers) at sufficiently high force magnitudes [2–5]. There have been observations that teams of kinesin motors can pull membrane tubes out of a vesicle [6]. This raises the question as to whether we should consider the alteration in vesicle shape due to tether formation in our computational model. But the previous analytical works [2, 3] have shown that the force magnitude needs to be greater than a critical value, 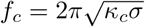 to pull a nanotube out of a membrane with bending rigidity *κ*_*c*_ and surface tension *σ*. For typical values, *k*_*c*_ = 20*k*_*B*_*T* = 10^−19^ J [5], *σ* = 10^−5^ Nm^−1^ [2, 7] the critical force is *f*_*c*_ = 8.8 pN which is higher than the stall force of kinesin (7 pN). This critical value that we computed is lower than the estimated value in at least one other study [2, 6]. We can see from the force distributions that we get from our simulations that the typical single motor force values are much lower than this critical value. Hence we have neglected the tether formation in our model.

## S2 Appendix

### Estimation of relevant timescales

#### *τ*_*bind*_ and *τ*_*off*_

*τ*_*bind*_ is the mean time taken for any one motor out of *N* motors of lipid cargo to bind to the microtubule. If a single motor takes *τ*_*ad*_ to bind, then if we have *N* of them, mean time for any one of the motor to bind is

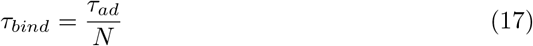

*τ*_*ad*_ = 1*/π*_*ad*_. We have measured the value of *π*_*ad*_ in Fig. 2(g) of main text for a lipid cargo of radius 250 nm and fluidity *D* = 1 *μm*^2^*s*^−1^. It is about 0.12 s^−1^. So *τ*_*ad*_ = 8.33 s.

*τ*_*off*_ is the time taken for a given bound motor to unbind. It is estimated as 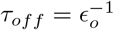, where *ϵ*_*o*_ is the off-rate of an unloaded kinesin motor (bare off-rate). Unloaded motor off-rate is a function of ATP concentration. At high ATP concentration of 2 mM, *ϵ*_*o*_ is measured to be 0.79 s^−1^ [1, 2]. Based on previous studies [3–5], we assume that *ϵ*_*o*_ = *v*_*o*_*/d. d* is the run length of unloaded kinesin motor which is found to be independent of ATP concentration. These information along with Michaelis-Menten equation provided in Eq. 10 of main text enable us to calculate unloaded velocity (*v*_*o*_), unloaded off-rate (*ϵ*_*o*_) and unbinding time (*τ*_*off*_) of a kinesin-1 motor at different ATP concentrations. Please refer to Table 1 below for numerical values.

#### Conditional on-rate

In Figs. 2(f-g), we showed that the binding rate of a motor increases as the number of bound motors increases. However, it might be challenging to measure this rate directly in experiments because the *n*-bound state is not stationary but can decay to *n* − 1 state by losing a motor. But one might be able to experimentally measure the number of bound motors as a function of time and one can obtain the *conditional on-rate per motor* from this data. We define the conditional on-rate per motor (S9 Fig.) as the rate of transitioning from *n*-bound state to *n* + 1-bound state divided by the total number of unbound motors in that state, *N* − *n*. A *n*-bound state can transition to a *n* + 1-bound state if any one of the *N* − *n* free motors binds to the microtubule. As mentioned earlier *n*-bound state can also decay to *n* − 1-bound state if one of the motors unbind. So we are looking at the rate with which *n* goes to *n* + 1 gated by *n* going to *n* − 1. To obtain this conditional on-rate from the number of bound motors as a function of time, we just have to filter all the *n* to *n* + 1 transitions in this time series, measure mean rate for such transitions and divide by *N* − *n*.

In addition to being straightforward to measure from trajectory data, it is also easy to find an analytical estimation of this conditional on-rate of a motor if we know the single motor on-rate and the single motor off-rate. Let 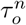 be the mean time for the decay of the n-bound state. Assuming that the motors work independently, we can write, 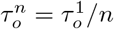. Let 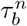 be the mean time for gaining 1 more bound motor. Since there will be *N* − *n* free motors when *n* motors are bound and these motors bind independently, 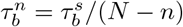 where 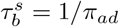 is the binding time of a single motor.

Mean time to go from *n* state to *n* + 1 state before *n* state decays to *n* − 1 state is

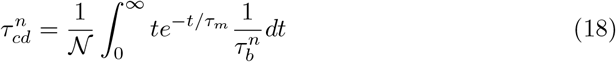

where

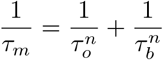

𝒩 is the normalization factor given by

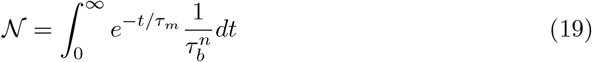

On simplification

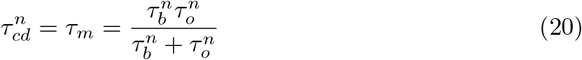

The conditional binding time per motor is 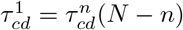. Conditional on-rate per motor is 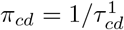.

We note that as *n* increases, the decay rate of *n*-bound state also increases. This implies the conditional on-rate per motor should increase as a function of the number of bound motors, just because of the increase in the decay rate (or decrease in the gating time) even if the binding rate of motor *π*_*ad*_ doesn’t change.

But we have seen in Fig. 2(g) that *π*_*ad*_ is not a constant value but is a function of n. We were curious to know how much difference will the change in *π*_*ad*_ make on the conditional on-rate as a function of n. So we plotted the estimated conditional on-rate taking *π*_*ad*_(*n*) (data from Fig. 2(g)) and compared with conditional on-rate with *π*_*ad*_ a constant (equal to *π*_*ad*_(1)) along with the measured value from simulations (S9 Fig.). However, there is not much difference between *π*_*ad*_(*n*) and *π*_*ad*_(1) case indicating that it might be challenging to observe this in experiments. The conditional on-rate measured from simulations agree well with the analytical estimations confirming the consistency between simulation measurements and analytical calculations.

## S3 Appendix

### Analytical estimation of run length for a rigid cargo

Since motors, are not free to move on a rigid cargo we approximate the run length as a weighted average of run lengths for the different number of motors in the access region [1]. The probability of locating exactly *l* motors in the access region when *N* motors are present on the cargo is

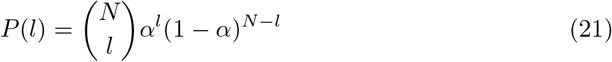

*α* is the ratio of the access region to the total surface area of the cargo. The run length of the rigid cargo is estimated as

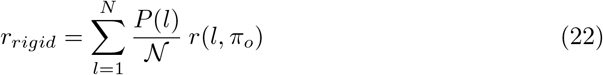

where 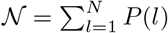 and *r*(*N*_*a*_, *π*_*ad*_) is given by [23]

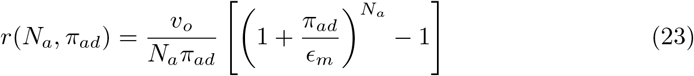

## S4 Appendix

### Cargo runlength can be different even though the average number of motors are the same

We observe in Fig. S11 that the average number of motors for N = 4 Low ATP (4.9 *μ*M) and N = 16 High ATP (2 mM) are approximately equal for lipid cargoes (*D* = 1 *μ*m^2^s^−1^). However, the runlength for the former case is higher than the latter. This is surprising since intuitively one would expect runlength to be the same when the average number of motors is the same.

We attribute this difference to the higher tendency to accumulate motors in the case of N = 4 Low ATP than N = 16 High ATP. A cargo run stops when the last bound motor unbinds. The likelihood for a new motor to bind before this last motor unbinds determines how far the cargo travels on the MT. Let us look at the ratio of the rate to bind one more motor to the unbinding rate for the case when there is just one motor bound, i.e.,

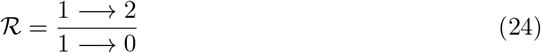

Calculated value of effective binding rate for a given motor in lipid cargo with *D* = 1 *μ*m^2^s^−1^ is approximately 0.12 s^−1^. So the rate for the process 1→2 is 0.12(N-1) s^−1^. The approximate unbinding rate, i.e., the rate 1→0 is 0.79 s^−1^ for High ATP and 0.079 s^−1^ for Low ATP (The no load off-rate of a single kinesin motor).

a. For N = 4 Low ATP (4.9 *μ*M)

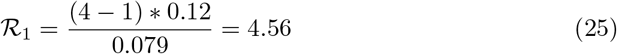
b. For N = 16 High ATP (2 mM)

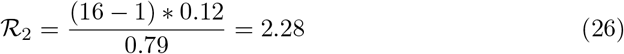

So the accumulation tendency in case (a) is higher than in case (b). Assume during a cargo run, the bound motor number becomes 1, then case (b) is more likely to fall off compared to case (a). However the average number of bound motors turns out to be the same because of the difference in upper bound, the maximum number of bound motors possible in case (a) is 4 whereas in case (b) it is 16.

